# A Synthesis of the Many Errors and Learning Processes of Visuomotor Adaptation

**DOI:** 10.1101/2021.03.14.435278

**Authors:** J. Ryan Morehead, Jean-Jacques Orban de Xivry

## Abstract

Visuomotor adaptation has one of the oldest experimental histories in psychology and neuroscience, yet its precise nature has always been a topic of debate. Here we offer a survey and synthesis of recent work on visuomotor adaptation that we hope will prove illuminating for this ongoing dialogue. We discuss three types of error signals that drive learning in adaptation tasks: task performance error, sensory prediction-error, and a binary target hitting error. Each of these errors has been shown to drive distinct learning processes. Namely, both target hitting errors and putative sensory prediction-errors drive an implicit change in visuomotor maps, while task performance error drives learning of explicit strategy use and non-motor decision-making. Each of these learning processes contributes to the overall learning that takes place in visuomotor adaptation tasks, and although the learning processes and error signals are independent, they interact in a complex manner. We outline many task contexts where the operation of these processes is counter-intuitive and offer general guidelines for their control, measurement and interpretation. We believe this new framework unifies several disparate threads of research in sensorimotor adaptation that often seem in conflict. We conclude by explaining how this more nuanced understanding of errors and learning processes could lend itself to the analysis of other types of sensorimotor adaptation, of motor skill learning, of the neural processing underlying sensorimotor adaptation in humans, of animal models and of brain computer interfaces.

## Introduction

The experimental study of sensorimotor adaptation has a history that predates the establishment of neuroscience and psychology as formal fields of study (von Helmholtz 1865), yet neither its psychological processes nor neural circuit mechanisms are fully understood. Some blame for this lies in the historical lack of agreement on the exact definition of adaptation and its constituent parts (Franklin and Wolpert 2011; Haith and Krakauer 2013; Held and Freedman 1963; Kornheiser 1976; Redding et al. 2005; Shadmehr et al. 2010; Shadmehr and Krakauer 2008; Wolpert et al. 2011). Here we broadly define sensorimotor adaptation as what takes place when a well-learned skill is modified to fit new conditions, such as when a carpenter borrows a hammer that is a different size than his own (James 1891). This review focuses on a narrower definition of adaptation as a motor learning process that alters future behavior in response to a discrepancy between actual sensory feedback and the sensory feedback that was expected to occur because of the motor command. When our carpenter uses the borrowed hammer, its different dynamics mean that his usual swing will result in unexpected tactile feedback in his hand, along with different visual feedback and potentially a different ultimate outcome for his intended goal. The motor system will automatically adjust itself in this case, tweaking the existing skill until the expected relationship between movements and feedback is re-established, usually reinstating adequate performance.

Adaptation is often studied in the laboratory through perturbation of limb motion during a movement, or by perturbation of the sensory feedback caused by the movement. There are many different examples of adaptation tasks, such as saccades and smooth pursuit (Gonshor and Jones 1976; McLaughlin and Webster 1967; Optican et al. 1985), disruption of walking gait (Dietz et al. 1994; Forssberg et al. 1980; Thelen et al. 1987), or reaching with external forces applied to the arm (Bock 1993; Fisk et al. 1993; Lackner and DiZio 1994; Sainburg et al. 1999; Shadmehr and Mussa-Ivaldi 1994). This review’s primary focus is visuomotor adaptation of manual reaching, as recent experiments have revealed much about the nature of this learning process. Visuomotor adaptation was originally induced by having participants view their arm through displacing prisms (Held and Hein 1958; von Helmholtz 1865), but can also be induced with completely artificial visual feedback mediated by an oscilloscope (Held et al. 1966), and more recently with computer monitors (Cunningham and Vardi 1990) or immersive virtual reality headsets (Anglin et al. 2017). We focus on visuomotor adaptation with the expectation that many insights gleaned from our analysis of this form of adaptation will generalize to other subtypes of sensorimotor adaptation and motor skill learning.

Historically, many adaptation studies focused on the locus of learning, specifically whether it was the senses, the motor commands, or the connection between these two domains that changed (Efstathiou et al. 1967; Welch 1969). This early work showed that vision, proprioception, and the relationship between the body and extrinsic space can all be altered in response to perturbed visual feedback. However, whenever visual feedback is perturbed, there is always a corresponding change in behavior that cannot be attributed to a retuning of the senses, meaning that its locus must lie either in the mapping between the senses and motor commands, or directly in the alteration of motor commands (see Kornheiser 1976; Redding et al. 2005 for thorough reviews of this topic). This “central” change to the visuomotor mapping is the specific focus of our review of visuomotor adaptation. Several studies suggest that this adaptation alters the displacement vector computed from the difference in the hand and target locations in extrinsic space, specifically altering the movement vector endpoint (Krakauer et al. 2000; Wang and Sainburg 2005; Wu and Smith 2013). Visuomotor adaptation can therefore be seen to primarily alter the step of the motor control hierarchy that transforms movement vectors computed in extrinsic space into intrinsic joint space coordinates, rather than the motor system as a whole (specifically, visuomotor adaption does not alter of the mapping between joint angle changes and the muscle activation required to change joint angles).

This consensus around what was primarily altered in adaptation inspired models of motor learning which assume that adaptation occurred via the modification of an internal inverse model (Atkeson 1989; Gomi and Kawato 1992; Hadjiosif et al. 2021; Raibert 1978; Shadmehr and Krakauer 2008). An internal inverse model takes the current state and a desired sensory state, or trajectory of states, as input and outputs motor commands. Many current models have a slightly different framework, positing that motor commands are generated using optimal feedback control, where motor commands are generated by a control policy that minimizes a loss function composed of the explicit desired sensory state and other factors like metabolic cost, pain and reward (Crevecoeur et al. 2014; Diedrichsen et al. 2009; Friston 2011; Kim et al. 2021; Pruszynski and Scott 2012; Scott 2004; Shadmehr and Krakauer 2008; Todorov 2004; Todorov and Jordan 2002; Wong et al. 2015). In this framework, some researchers suggested that adaptation occurs via the adaptation of the internal forward model (Crevecoeur et al. 2020a, 2020b; Izawa et al. 2008). An internal forward model takes the generated motor commands as input and outputs the predicted sensory consequences of the movement. This internal forward model is taken into account by the control policy during planning. In the optimal feedback control framework, adaptation of the internal forward model leads directly to a change in generated motor commands because it modifies the state estimate (Diedrichsen et al. 2009). Notwithstanding these subtle differences between frameworks, a consensus view is that implicit sensorimotor adaptation (specifically the adaptation of the internal inverse and/or forward models) alters the outputs of the control policy.

It is important to disambiguate the above learning, which represents implicit adaptation, from the use of an explicit strategy to compensate for imposed perturbations. A cognitive strategy can encompass many things. In this context we are referring to the explicit decision to re-aim, or change the spatial goal of a reach, so that the participant intentionally plans to move their hand to a location that is distinct from the location of the target. Disambiguation between explicit and implicit adaptation requires specific experimental procedures (Hadjiosif and Krakauer 2020; Maresch et al. 2020a), as the two types of learning result in the same behavior: a single reach that has been adapted to move 15° off midline is kinematically identical to an un-adapted reach that is aimed 15° off midline. Although people tend to fixate the location where they are aiming, this is not necessary (de Brouwer et al. 2018; Rand and Rentsch 2015), meaning that there is not necessarily an external behavioral marker of implicitly or explicitly adapted movements. Similarly, in other movement contexts, the aiming point and the target are not always identical. For instance, while preparing to fire a single arrow, an archer may notice that the wind is blowing to the left and deliberately decide to aim her shot to the right of the bull’s eye to compensate. Such a strategy does not reflect a change in the control policy, and it would be inappropriate to characterize it as implicit adaptation to the wind.

We give these examples to demonstrate that the use of an explicit aiming strategy and implicit adaptation could lead to the same change in measured behavior, despite vastly different underlying mechanisms in terms of the motor control hierarchy, neural loci and computational processes (McDougle et al. 2016). In early work, a minority of researchers sought to dissociate the explicit and implicit components of visuomotor adaptation (Weiner et al. 1983; Welch 1969). Today measuring or experimentally controlling the use of strategies during adaptation tasks is becoming a standard in the field (Butcher et al. 2017; Haith et al. 2015; Hegele and Heuer 2010; Heuer et al. 2011; Heuer and Hegele 2011, 2014; Leow et al. 2017, 2018, 2020; Mazzoni and Krakauer 2006; McDougle et al. 2015, 2017; Miyamoto et al. 2020; Morehead et al. 2015, 2017; Rand and Heuer 2020; Ruttle et al. 2020; Schween and Hegele 2017, 2017; Taylor et al. 2014; Tsay et al. 2020b; Vandevoorde and Orban de Xivry 2019, 2020a). It is also increasingly recognized that there is a graded boundary between implicit and explicit behavior, which can complicate the differentiation these processes (Hadjiosif and Krakauer 2020; Maresch et al. 2020a).

This focus on the implicit/explicit dichotomy led to the observation that these processes may be driven by distinct error signals (Shadmehr et al. 2010; Taylor et al. 2014), with sensory prediction-error and performance error being of particular interest (see below for a definition of these terms, Jordan and Rumelhart 1992). Indeed, recent experiments designed to measure behavioral responses to distinct errors have revealed that there are likely more than two distinct learning processes contributing to visuomotor adaptation. Our primary purpose here is to review this disparate experimental work to describe a coherent and holistic view of reliable data about how visuomotor adaptation occurs in the natural world. Moreover, we believe such a framework will be useful for a diverse group of scientists in the design and interpretation of motor learning and skill acquisition experiments, for both behavior and neurophysiological measures.

## Visuomotor adaptation is driven by different error signals

Many visuomotor adaptation studies are performed in the center-out reach paradigm (Georgopoulos et al. 1981). This type of task features planar reaches made from a central start position to targets arranged in a circle around the starting point (Fig. 1a, baseline). Visual feedback of their hand position is represented via a cursor on the screen, while direct vision of the hand is typically occluded. In these tasks, people are asked to either to bring their hand to rest within the target or to pass their hand through the target without stopping. Bringing the hand to rest within the target location typically takes more time, and success in this type of reach often requires online correction of the movement. Movements where the hand briefly passes through the target can be made quickly enough (150-200ms) to ensure that online feedback corrections from sensory feedback are minimized or precluded entirely (Elliott et al. 2010; Saunders and Knill 2003). Visual feedback of the hand position is typically presented to provide information on the accuracy of a participant’s movements, either online during the reach or at designated points during the movement, such as the endpoint. This visuospatial feedback of hand position is often supplemented with additional information on movement duration and spatial accuracy via auditory tones, text displays, or changes in the color of objects on the display.

**Figure 1.**
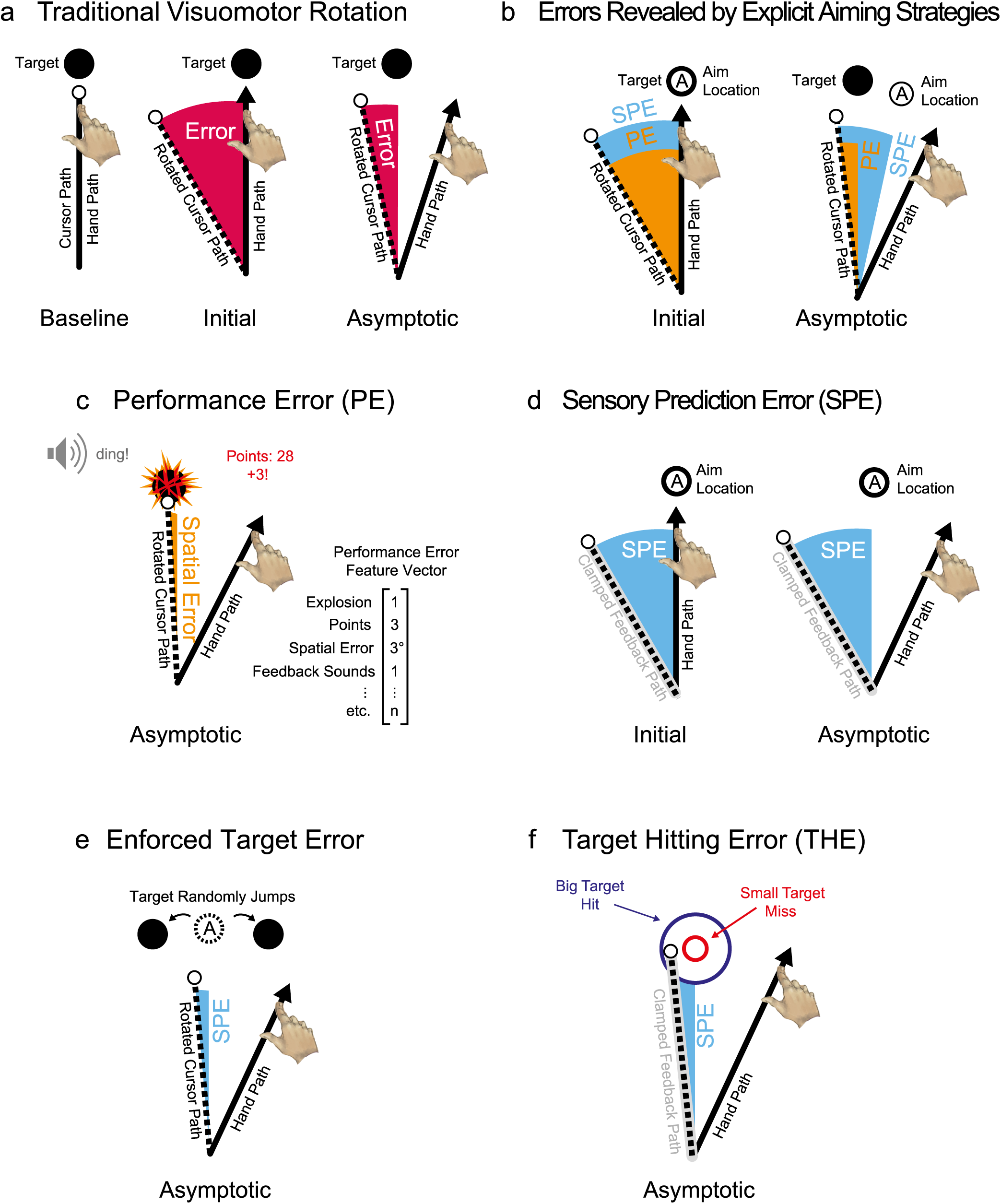
Errors in Visuomotor Adaptation Task Contexts. A) A traditional visuomotor rotation task. The hand of the participant, hidden from view, moves from a start position to the target (black circle). During baseline, the feedback cursor (white circle), shows the actual location of the hand. When feedback is perturbed, the position of the cursor is rotated around the start position, so that it moves in a different direction from the hand when a movement is made away from the center. Later in learning, the participant has altered the direction of movement to counteract the rotation of feedback, bringing the cursor closer to the target (asymptotic learning). The difference between the cursor and target angle (in pink), is the error signal driving learning in most models of motor learning. We refer to this error as the composite error. B) The change in movement direction that counteracts the cursor rotation can be made implicitly or explicitly. Experiments that measure or control explicit aiming strategies reveal that these two types of correction are driven by at least two error signals: explicit strategies by performance error and implicit adaptation by sensory prediction-error. The aiming direction is represented by the A symbol within a circle on the right panel. The task performance error (in orange) is the difference in angle between the target and the cursor. The sensory prediction-error (cyan) is the difference between the aiming direction and the direction of the cursor. C) Performance error in visuomotor adaptation is more than a visuospatial error vector between the cursor and target. On a hit it may include any signal that conveys information about task success, including a visual “explosion” of the target, a pleasant sound and/or numerical points provided after the movement. For a miss, the target may not explode, the sound may be unpleasant and/or points could be withheld or subtracted. These signals of success may accompany or take the place of visual feedback of position. Performance error may best be thought of as a feature vector made up of many disparate elements from sensory and cognitive modalities. D) Sensory prediction-errors are ostensibly generated when sensory feedback is compared to that predicted from the motor command by a forward model. Here we show a visual signal that is thought to generate such an error, isolated from performance errors with the task-irrelevant clamped feedback method. Here the cursor always moves at a fixed angle that is offset from the target, and participants are informed that they cannot control the direction of this feedback. Moreover, they are instructed not to try to control the feedback and to instead ignore it while always moving their unseen hand directly to the target. This manipulation induces implicit adaptation without any changes in aim or performance error throughout the task. Notably, because the clamped feedback angle is fixed and aim is always at the target, the putative sensory prediction-error also remains unchanged throughout the task. E) Another way to isolate adaptation to a sensory prediction-error is to randomly change the sign of performance error on every trial by jumping the reach target out of the way during the second half of movement. This allows trial-to-trial measurement of the same implicit adaptation as task-irrelevant clamped feedback without giving participants information on the nature of the perturbation or providing specific strategies to ignore or aim to a location. There should be no net change in aim in this task over the course of a learning block or within a group of participants as the sign of the jumps is random. The target can also be jumped directly into the path of the cursor so that the movement is always successful. F) The angular deviation of the feedback is fixed with the task-irrelevant feedback method, and at small offsets this may intersect the target if it is large (blue circle, Big target hit) or miss the target when it is small (red circle, small target miss). Along with the target jumping method on the left, this method revealed that there is a third signal, a seemingly binary target hitting error, that modulates implicit adaptation even when it is irrelevant to task performance.

### Composite error (CE)

In the visuomotor rotation task (Fig. 1A, initial and asymptotic), feedback is perturbed by applying a rotation to the position of the hand feedback cursor relative to the start position. This only alters the angle of the feedback, independent of the radial position, meaning the magnitude and speed of the feedback remains unaltered and statistically independent of the perturbation (Krakauer et al. 2000; Pine et al. 1996). Such rotation introduces an error (Fig. 1A), or a difference between what the participant planned to do and what occurred. This error may result in a failure to hit the target entirely, or the task may require the error to be corrected online during the trial. In either case, the imposed error drives adaptation of the reach behavior on subsequent trials, which decays once the perturbation is removed (Tseng et al. 2007). Although the discrepant visual feedback was the only error introduced directly, this usually causes a cascade of other errors. For instance, if the visual feedback cursor did not reach the target in time, the trial may be considered less rewarding by either the participant or the experimental software (registered via sounds for failure, a lack of points awarded, etc.). Internal to the participant, the moment to moment sensory feedback that is predicted for each movement by a forward model will be discrepant. Taken together, these error signals form a composite error that drives adaptation of reaching behavior. This composite error is used by almost all models of motor adaptation until now, as most experimenters did not have a way to differentiate between error types. Recent research has shown that this error can be broken down into constituent parts, and that these unique facets of error drive different learning processes that cause changes in behavior during adaptation tasks. Below, we detail the different constituent error signals that have been identified from the composite error that is introduced by visuomotor perturbations.

### Performance Error (PE)

Performance error is a term for any feedback that conveys information about success in any undertaking. We want to emphasize that performance error is specific to each task and task environment. In natural settings, the task is determined entirely by the organism, as the animal defines its own goals via instinct or reason and finds sensory feedback that is predictive of success (Skinner 1981; Tinbergen 1951). In the laboratory, the task is often arbitrary, with the experimenter relating an understanding of the task and what constitutes success via verbal instruction, imitation, and interactive feedback from the task itself. Participants further refine their understanding of the task and its signals for performance error through practice and questions of the experimenter.

In a reach adaptation context, the primary performance error metric is the difference between the visuospatial feedback of target position and the hand (Fig. 1C, spatial error). This feedback can be given during or after a movement is complete and may reflect the entire movement or only specific points in the movement. All of these constitute a graded, or scalar, form of feedback that could be reported in one, two or three dimensions. Importantly, some of the visuospatial feedback may be superfluous to the participant’s task (e.g. “hit the target”), or all of it could be important (e.g. “move straight and hit the target”). Additionally, a very common task success metric is movement time, which may be reported to the participant numerically, graphically (a number line or bar graph), or with visual or auditory categorical metrics (“too slow”, “too fast”). Depending on the task, this space and time feedback may be supplemented by categorical or graded feedback such as pleasant or aversive sounds, “explosions” of the target, points that increment or decrease, money that is earned or lost (Fig. 1C), and primary rewards such as juice or food (in the case of animal experiments). These signals are all sources of performance error. Therefore, performance error should be thought of as a multidimensional error signal gathering all the relevant aspects of the task performance. Put differently, performance error can be thought of as a feature vector of the many factors that may themselves be scalar, categorical, or binary (Fig. 1C).

Participants can use performance error feedback, however coarse, to reinstate good performance in response to a perturbation (Butcher et al. 2017; Nikooyan and Ahmed 2015). Such learning can take place gradually or in abrupt steps, and can appear exponential—especially if averaged across participants (Gallistel et al. 2004). The evidence suggests that learning from performance error does not recalibrate internal models, instead it arises primarily from alterations in explicit aiming strategies (van Beers 2009; Wong et al. 2019). This observation is controversial, as it is a long-held view that adaptation occurs in response to performance error (Atkeson 1989; Jordan and Rumelhart 1992). This view was tenable because most studies unknowingly measured a composite of several error types rather than performance error in isolation.

Relevant to performance error is the difference between experienced and predicted reward is known as reward prediction-error (Montague et al. 1996). Critically, this is not a primary signal in the environment, but a signal calculated internally by an organism (or simulated by a model). This expectation of reward can be conditioned either passively or actively through classical or instrumental conditioning, respectively. Generation of this error signal is useful for maintaining a well-calibrated valuation of states, stimuli, actions and outcomes (Schultz 2016). It can therefore be seen to indirectly affect learning in adaptation tasks as a variable intrinsic to non-motor decision making. We assume that reward prediction-error is being calculated relative to both graded and punctate performance errors (including primary rewards and punishments, when present) that are experienced on each trial (Fig. 1F). We therefore do not discuss it in further detail, other than to point out that it is intrinsically involved in an organism’s internal assessment of its own performance and economic utility in general.

### Sensory Prediction-Error (SPE)

There is good agreement that the motor system feeds corollary discharge of the descending motor commands to an internal forward model so that the future state of limbs can be estimated during movement (Franklin and Wolpert 2011; Shadmehr et al. 2010; Shadmehr and Krakauer 2008; Wolpert et al. 2011; Wolpert and Kawato 1998; Wolpert and Miall 1996; Wong and Shelhamer 2010). It is hypothesized that this sensory prediction is compared to the actual sensory feedback to form the sensory prediction-error, which has long been thought to play a role in adaptation (Jordan and Rumelhart 1992). Importantly, sensory predictions and sensory prediction-errors are generated internally by the organism and can only be indirectly measured or manipulated, barring direct stimulation or recording of neurons in the circuit (Pasalar et al. 2006; Requarth and Sawtell 2014). In visuomotor adaptation experiments, the perturbation of sensory feedback ostensibly induces a sensory prediction-error (Fig. 1D), which results from the difference between the sensory feedback predicted from the motor command (here, a spatially and temporally specific motion of a visual cursor) and the actual sensory feedback (motion of the feedback cursor). In Figure 1D, we show this putative sensory prediction-error in the context of the task-irrelevant clamped feedback task, where participants are informed that the cursor will travel along a specific pre-determined path that is angularly offset from the target and, importantly, that this feedback does not show the position of their hand. Moreover, participants are told that their task is to ignore this motion stimulus and move their unseen hand directly to the location of the target. Despite instructions to ignore this feedback, and an explicit understanding that this feedback is not related to their performance in the task, participants show a robust implicit adaptation that is indistinguishable from implicit adaptation observed in response to conventional visuomotor rotations (Avraham et al. 2021; Kim et al. 2018, 2019; Morehead et al. 2017; Poh et al. 2021; Tsay et al. 2020b, 2020c, 2020a, 2021; Vandevoorde and Orban de Xivry 2019, 2020b). In these other tasks, rotated feedback has been shown to drive implicit adaptation independent of performance error (Kim et al. 2018; Leow et al. 2018; Mazzoni and Krakauer 2006; Miyamoto et al. 2020; Morehead et al. 2017; Taylor and Ivry 2011; Vandevoorde and Orban de Xivry 2019; Wolpert et al. 1995). For both task-irrelevant clamped feedback and relevant perturbed feedback, this adaptation presumably occurs because it induces a sensory prediction-error.

In classical visuomotor rotations, sensory prediction-errors and performance errors are conflated the early stage of adaptation, where the perturbed feedback should be considered as a composite error signal (initial adaptation in Fig. 1A). If the participant changes their aim from the visual target’s location, their performance error and sensory prediction-error signals will differ from this point in the adaptation period. It is therefore easier to dissociate these errors in tasks where explicit aim is measured (Fig. 1B). Here performance error is represented by the difference between the direction of the cursor’s motion and the target angle. Sensory prediction-error, on the other hand, corresponds to the difference between the angle of aim and the visual feedback’s direction of motion. It should be noted that sensory prediction-error is a hypothesized signal has not been directly observed in sensorimotor adaptation experiments despite strong theoretical foundations.

### Target Hitting Error (THE)

Target hitting error is a binary error signal that modulates behavior irrespective of the explicit task context. We introduce this term because two independent studies have shown that vision of a cursor intersecting the target of a reach attenuates implicit visuomotor adaptation (Kim et al. 2019; Leow et al. 2018). In one of these studies (Fig. 1D), during the adaptation to a visuomotor rotation, the target of the reach jumped into or away from the path of the cursor. In both types of target jump, the typical relationship between behavior and performance error was eliminated, as the participant was either always successful or unsuccessful. Despite this decoupling of performance error, the authors observed that the adaptation was larger when the target jumped away from the cursor path than when it jumped into the cursor path (Leow et al. 2018). In the other study (Fig. 1E), the researchers used the task irrelevant clamped feedback protocol described above (Fig. 1C) but varied the width of the target (Kim et al. 2019). In the hit condition, the target was large, and the cursor reached the target, while in the miss condition the target was smaller and the cursor always missed the target. Here again, adaptation was larger in the miss condition than in the hit condition. In both sets of experiments, the participants had no control over whether the cursor hit the target, and in one case were explicitly told not to attempt to control this element of the task. Nonetheless, in both cases the task-irrelevant motion of the cursor through the target attenuated the amount of implicit adaptation. It should be noted that both studies showed evidence for a binary, or categorical, target hitting error; there is no published evidence that this signal varies parametrically with the distance of the center of the cursor and target.

## Adaptation consists of distinct processes driven by different errors

The dynamics of behavioral changes during sensorimotor adaptation has been interpreted in many different ways, depending largely interests of researchers (Kornheiser 1976; Redding et al. 2005). Here we detail several influential theoretical and task-level descriptions of adaptation, some of which are mutually exclusive. Our later synthesis will seek to explain how the combined behavior of several independent learning processes can explain the behavior observed in each of these contexts.

Adaptation to a visuomotor rotation is typically considered as an error learning process where the hand direction on a given movement is determined by a visuomotor mapping from the perceived target direction to a motor command that will move the hand in the same direction. For a movement generated towards a given target direction, the hand’s direction of motion depends on the internal state (*X_k_*) of this visuomotor mapping and execution noise (noise).

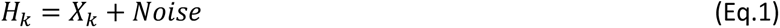

This visuomotor map is assumed to be at a well-calibrated steady state in the absence of external perturbations, making only small corrections for execution errors. If an external perturbation is introduced, the system will retune its mapping weights, departing from its steady state by a fixed fraction of the error (*e*) size, parameterized as a learning rate (*b*). Early models only included the error and learning rate for the state update on every trial (Atkeson et al. 1988; Donchin et al. 2003; Raibert 1978; Thoroughman and Shadmehr 2000), but later models introduced a forgetting term (Smith et al. 2006). This forgetting factor (*a*) causes the internal state of adaptation to decay by a fixed percentage between trials. It accounts for two observations, that the adapted memory decays in the absence of feedback (van der Kooij et al. 2015, 2016) and that adaptation is rarely complete, typically leaving an asymptote that is 5% or more short of the imposed perturbation. The following state-space equation illustrates this mechanism. The state of adaptation on the next trial is a function of the amount of adaptation on the previous trials *X_k_*_−1_ weighted by a forgetting factor (*a*) summed with the error observed during the previous movement *e_k_*_−1_ weighted by a learning rate (*b*):

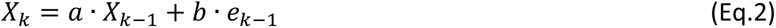

This simple state-space model describes the dynamics of the adaptation process using the composite error (*e_k_*_−1_) as a signal for learning. Such a model shows a remarkable flexibility, learning to compensate for the perturbations in a large variety of situations (Cheng and Sabes 2006; Thoroughman and Shadmehr 2000) (Fig. 2A). Nonetheless, it suffers from some limitations.

**Figure 2:**
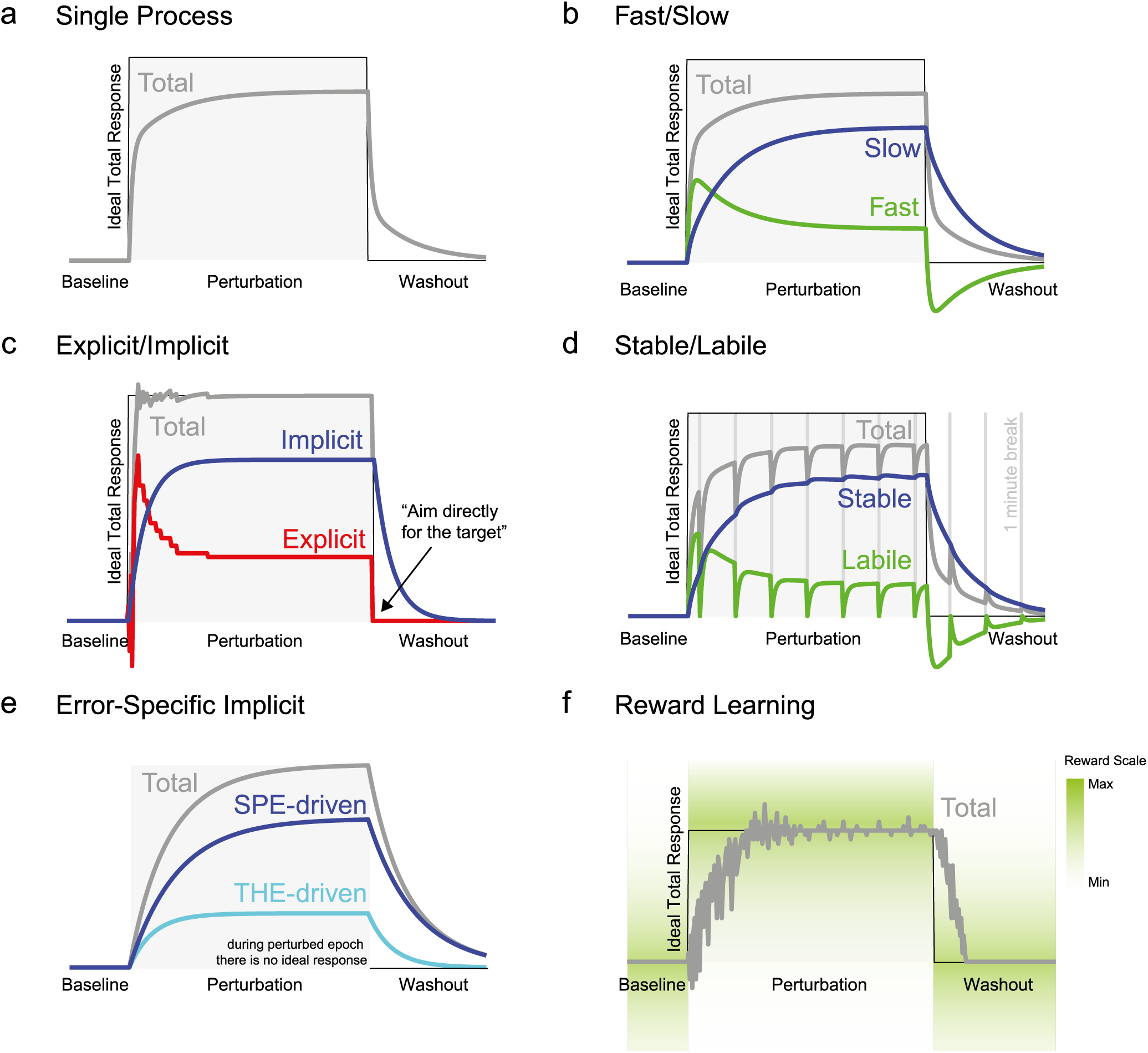
Time course of distinct processes hypothesized to underlie visuomotor adaptation. A) Single process. When a visuomotor rotation is introduced at the end of baseline, the motor system adapts the direction of the hand’s motion (grey curve) towards the ideal total adaptative response (black trace), which is the opposite of the feedback rotation angle. This change in hand direction over the course of trials represents the total adaptive response. Upon removal of the perturbation (washout period), the adaptation response returns gradually to baseline levels. B) Fast and slow processes. The total adaptation response can be decomposed into a fast process that learns fast but forgets rapidly (green curve) and a slow process that evolves slowly but has good retention properties (blue curve). During washout the fast process takes negative values in order to bring the total adaptation curve to baseline levels as fast as possible. C) Explicit and implicit processes. The total adaptative response can be decomposed into an explicit and an implicit process (red and blue curves, respectively). At the start of the perturbation period, the explicit process is extremely variable and can take positive and negative values as the participant is experiments with different strategies to cope with the perturbation. In contrast, the implicit process evolves more slowly and monotonically. During the washout, participants are instructed to aim their movement directly to the target (often without feedback) so that the implicit process can be measured directly. The implicit curve during the rotation is not measured, but estimated by subtracting reported aim from the measured reach direction. C) Labile and stable processes. The processes underlying motor adaptation can be defined as a function of their resistance to the passage of time. In this case, interspersing 1-minute breaks throughout the experiment allows one to look at the effect of time on the adaptation response. Two processes emerge from this manipulation: a stable process whose response does not decay during one-minute breaks and a labile process that decays substantially during these breaks. The labile process appears to learn to both learn and forget faster than the stable process. E) Error-specific implicit process. Implicit adaptation to a putative sensory prediction-error differs depending on whether the target was hit by the cursor or not. Specifically, there is less adaptation when the target is hit. Therefore, it has been hypothesized that the implicit adaptation response could be divided into a sensory prediction-error (SPE) driven and a target hitting error (THE) driven process (blue and cyan curves, respectively). F) Reward learning process. In the absence of any visuospatial feedback about the participant’s performance, the adaptative response can be guided by reward signals such as binary feedback (with gradual perturbation) or with graded feedback (e.g. points, a tone) that varies relative the distance from ideal behavior. In this case, learning is extremely variable but slowly evolves towards the ideal correction for the imposed perturbation.

### Fast-slow learning

Single process state-space models cannot account for the double exponential learning curves (initial fast learning period followed by slower learning period) that are commonly observed in adaptation nor can they account for spontaneous recovery of the motor memory and savings. To address these shortcomings, Smith *et al*. (2006) described sensorimotor adaptation as two learning processes operating in parallel: a fast process (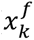, red line in Fig. 2B) that learns quickly (*b^f^*) but has poor retention (*a^f^*) and a slow process (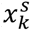, blue line in Fig. 2B) that learns slowly (*b^s^* < *b^f^*) but has good retention of its state from one trial to the next (*a^s^* > *a^f^*).

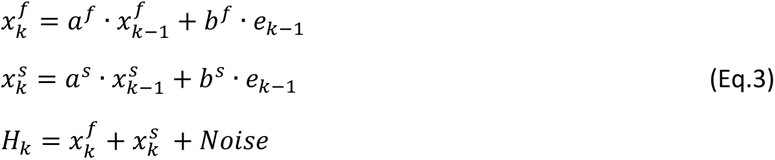

Both processes are driven by the same composite error signal (*e_k_*_−1_), and their learning is summed to affect behaviour with the addition of execution noise (blue line in Fig. 2B), although the processes operate independently. As represented on Fig. 2B, it is interesting to note that the proportional contribution of each process to the total adaptation varies over the course of training. This dual process model can explain double exponential learning curves, and, in limited contexts, the existence of spontaneous recovery and savings of motor memories (Smith et al. 2006; Zarahn et al. 2008). It can be further extended to account for more than two timescales (Kording et al. 2007).

### Explicit-Implicit learning

Sensorimotor adaptation has components that can be categorized based on whether they are accessible by explicit conscious awareness or not. Such distinctions are fundamental for understanding the neurophysiology of memory (Squire 2004). In the context of visuomotor rotation, the explicit process we refer to corresponds to a deliberate and conscious aiming strategy used by a participant to counteract the perturbation (Taylor et al. 2014; Welch 1969). For instance (Fig. 1B), if a cursor deviates towards the left because of a rotation, the participant might explicitly think “aim to the right” of the actual target in order to compensate for the effect of the rotation on her motor performance. This process is explicit because the participant consciously plans to move to a different spatial location than the visual target location. This explicit decision is under volitional control, meaning that the participant may change the location of their aim at will.

There are different ways to directly measure explicit aiming in center-out reach tasks. The most straightforward is to have participants report the location of their aim for each reach. Experimenters have had participants report their aim verbally, either by placing landmarks on the screen that participants verbally reference, or by having participants call out imagined numbers on a clock face (Benson et al. 2011; Taylor et al. 2014). For instance, in the study of Taylor and colleagues (2014), numbers were presented on a circle around the starting position. The target corresponded to the number zero. Numbers increased with angular deviation in the clockwise direction and decreased (i.e. became increasingly negative) in the counter-clockwise direction. Before each movement, the participants were required to verbally report the number indicating where they intended their hand to move to (i.e. their “aim”). Aim can also be measured by allowing the participant to position a marker in the workspace with computer peripherals such as a keyboard (Miyamoto et al. 2020), or they can indicate the point of aim by pointing to a touch-screen with their other hand (Bond and Taylor 2017). Eye fixation can also be used as a noisy proxy for explicit aim, but it is not a reliable enough to supersede explicit reports (Bromberg et al. 2019; de Brouwer et al. 2018). Moreover, one can employ an aiming strategy at a location that is different from their gaze position (Rand and Rentsch 2015). The difference between these trial-to-trial reports of the intended reach location and the actual reach direction represent implicit adaptation and motor execution noise. When explicit aiming is measured with these techniques, it is typically found that this component of motor learning quickly reaches a high point in the early adaptation period, then slowly decreases to a smaller fixed value. The implicit component evolves more slowly, growing monotonically until it reaches an asymptotic state (Fig. 2C). Several studies have shown that the explicit component is largest during early adaptation to a fixed perturbation, and that it declines but does not entirely go away with extended exposure (McDougle et al. 2015; Morehead et al. 2015; Taylor et al. 2014). Similarly, restricting reaction time has been used to limit the amount of explicit adaptation (Haith et al. 2015; Leow et al. 2017) but this method suffers because explicit strategies can be cached with training and expressed at low preparation times (Huberdeau et al. 2019; McDougle and Taylor 2019). Importantly, this means that reaction time restrictions do not eliminate the use of explicit aiming strategies.

Rather than estimating implicit adaptation by subtracting explicit aim from behavior, the accumulated amount of implicit adaptation can be measured directly by giving participants strong instructions on where to aim and withholding feedback at points within a perturbation block or afterwards (Avraham et al. 2020; Hegele and Heuer 2010; Heuer et al. 2011; Heuer and Hegele 2014; Morehead et al. 2015, 2017; Taylor et al. 2014; Taylor and Ivry 2011; Vandevoorde and Orban de Xivry 2019, 2020a). For instance, in the washout period of Fig. 2C, we illustrate a scenario where participants are instructed to aim directly at the target following a block of perturbation trials. Here, explicit aim is directed to the target location (at 0°) and implicit adaptation causes the hand to deviate from the straight ahead path seen during baseline. While we illustrate this technique for a series of trials on Fig. 2C, this can be also applied for single trials interspersed throughout the adaptation block. In this case, the participants are cued that they should abandon aiming strategies on the next trial. This no feedback catch trial/aftereffect method can be employed in a wide variety of visuomotor adaptation contexts and has the advantage of not providing the participant with a specific strategy. However it has the disadvantage of disrupting the learning process, which introduces a decay of the adapted state, and subsequent learning cannot be considered naïve to externally suggested modifications in aim.

The idea that sensorimotor learning has explicit components is not new (Hendrickson and Schroeder 1941; Welch 1969). However there has been a great deal of interest in explicit adaptation since it was clearly shown to play a large, persistent, role in sensorimotor adaptation, and that it bears a striking similarity to theorized fast and slow adaptation processes (McDougle et al. 2015). Research on methods for the measurement and control of explicit and implicit learning is progressing rapidly (Hadjiosif and Krakauer 2020; Maresch et al. 2020b), offering many different options that cannot all be detailed here. Care should be taken in navigating this explicit/implicit dichotomy. One thing is clear, there are explicit and implicit learning components of adaptation across all forms of adaptation, independent of the type of perturbation. The measured magnitude of these processes may depend in part on the method used to measure them, and we know the proportion of explicit learning out of the total adaptation differs as a function of the perturbation and the experimental context (Bond and Taylor 2015; Schween et al. 2020). Importantly, throughout this review when we discuss explicit strategies, we are referencing strategies that are consciously and volitionally employed by the participant at the time of their use, not only upon reflection at a later point in time (Hadjiosif and Krakauer 2020).

### Labile-stable learning

Memories for sensorimotor adaptation also vary in their evolution over time. Some show a time-dependent decay within a minute, while others are relatively stable over hours or days. This is not particularly surprising, as it is common to observe that a portion of adaptation decays during breaks when no movements were made (Fig. 2D). The time course of this decay has been systematically investigated over minutes (Alhussein et al. 2019; Hosseini et al. 2017; Joiner et al. 2017; Joiner and Smith 2008; Sing et al. 2009; Zhou et al. 2017). This work suggests a simple dichotomy between the component that decays over time, dubbed temporally labile memory (green line on Fig. 2D), and the component that only decays when movements are made, the temporally stable memory (blue line on Fig. 2D).

### Learning from target “hits”

Two studies demonstrated that the intrinsic performance error directly influence the implicit adaptation process (Fig. 2E, Kim et al. 2019; Leow et al. 2018). Here will illustrate this process as observed in the task-irrelevant clamped feedback protocol described above (Fig. 1D) with the use of small and large targets (Fig. 1F). This work showed that the adaptation process did not reach the same plateau in the hit and miss conditions, even though the angular offset of the clamped feedback was identical. When the cursor entirely missed the target, the plateau of adaptation was larger as it resulted in the sum of the implicit adaptation (blue line in Fig. 2E) and of target hitting error process (cyan line in Fig. 2E). In the hit condition, this additional component was absent, and total adaptation remained at the level of the SPE-driven adaptation only. The presence of the intrinsic performance component is thus best demonstrated by contrasting a hit and a miss condition as done in both studies. In both cases, implicit adaptation was modulated by whether the cursor was seen to have hit the target, despite that event’s relevance to explicit performance error.

### Instrumental/Reward learning

A variety of studies show that reinforcement learning, or instrumental conditioning, can learn to compensate for a visuomotor perturbations (Butcher et al. 2017; Cashaback et al. 2017, 2019; Codol et al. 2018; Darshan et al. 2014; Holland et al. 2018; Izawa and Shadmehr 2011; Mastrigt et al. 2020; Palidis et al. 2019; Pekny et al. 2015; Therrien et al. 2015). For instance, in such studies, participants are required to reach to a target without visual feedback relating to their hand position during or after their movements. In binary feedback experiments of this type, the only external feedback participants receive is a hit or miss signal (e.g. the target changes color if you hit it). In versions of this task with graded (scalar) feedback, participants may see points or some other non-spatial representation of feedback that indicate the magnitude and/or direction of their reach accuracy relative to the target. To perturb this behavior, the region of the workspace that will trigger the feedback for a target hit is rotated by a fixed angular offset. This requires the participant to change their hand angle in order to receive the same reward feedback that they did under baseline conditions. In graded reward contexts (Fig. 2F), convergence to the new ideal behavior usually progresses on a similar time scale to traditional visuomotor rotations with full online visual feedback. If the feedback is binary, participants do not have any information about the direction of the perturbation and have to find the new reward region purely through exploration, which may result in them not finding the new solution (Manley et al. 2014).

This illustrates that any performance error learning problem, such as visuomotor rotation, can in principle be solved by reinforcement learning (Jordan and Rumelhart 1992). Importantly, though these distinct learning algorithms can eventually solve the same problems, there is a mechanistic distinction between the internal process used by unsupervised reinforcement learning and “error-driven” supervised learning processes that are thought to play a major role in implicit sensorimotor adaptation. Such a difference in the nature of the operant process will result in different trial-to-trial behavior during learning. Fundamentally, reinforcement learning algorithms maximize reward by sequentially exploring the action space via random sampling or hypothesis generation, to in turn exploit actions with better outcomes (Dayan and Daw 2008; Marr 1970). Conversely, supervised learning adjusts its actions relative to the error between the teaching signal and feedback from each trial to the next (Marr 1969; Raymond and Medina 2018). In sensorimotor adaptation, the teaching signal for supervised learning is thought to be the sensory prediction-error (Jordan and Rumelhart 1992; Shadmehr et al. 2010). The gross learning curves of these processes can appear similar, especially in the presence of execution noise, but their operation is distinct and should be distinguishable with the right experimental design (Cashaback et al. 2017).

There is little doubt that reinforcement learning plays a role in decisions about explicit strategy use in visuomotor adaptation (McDougle et al. 2016). It is unclear, however, the extent to which extrinsic reward and punishment affect implicit learning. Reward and punishment have been shown to modulate learning in adaptation tasks, with differential effects on retention (Cashaback et al. 2017; Darshan et al. 2014; Gajda et al. 2016; Galea et al. 2015; Izawa et al. 2012; Izawa and Shadmehr 2011; Nikooyan and Ahmed 2015). It has also been argued that rewards associated with specific actions bolster retention of motor memories (Galea et al. 2015; Huang et al. 2011; Shmuelof et al. 2012). However, many of the effects in these studies may arise entirely from explicit strategies (Codol et al. 2018), as studies on adaptation that involve operant learning have often not controlled for this factor.

## A non-motor stage for the control of visuomotor behavior

So far, we have discussed the existence of the numerous processes operating during sensorimotor adaptation, and the fact that these processes appear to be driven by distinct error signals. Here we offer a new framework that differs from previous attempts because it distinguishes different types of errors and articulates which learning processes are driven by these errors. We believe this synthesis can help explain many counter-intuitive phenomena that have accumulated in the motor learning literature in recent decades and that it can guide future studies on the neural basis of motor adaptation. This synthesis is needed because many of the points of view we outlined above were developed to explain the totality of behavior in sensorimotor adaptation, meaning that they are fundamentally at odds. In some cases, this has led to fierce controversies. For instance, the claim that explicit strategies constitute fast learning in sensorimotor adaptation pits multi-rate models (Smith et al. 2006) against explicit/implicit learning (Taylor et al. 2014) but these ideas are not actually mutually exclusive. It is likely that implicit adaptation has processes that learn at different rates, and an explicit strategy can change more rapidly than implicit learning but is also likely to be retained over 24 hours (Wilterson and Taylor 2019). There are also many cases where a failure to consider the different predictions of these frameworks can lead to a general misunderstanding of what behaviors mean in these tasks. We will explore some of these interactions further below.

To describe this new framework, we have first updated a traditional sensorimotor control and learning diagram, with the only substantial changes coming at the input stage. Classical control diagrams in the motor learning field start with a “desired” sensory feedback, or sensory goal (Atkeson 1989; Jordan and Rumelhart 1992; Raibert 1978). In previous models, the goal was transformed into motor commands via an internal inverse model (or optimal controller). In the latter case, the motor commands are computed in relation to the loss function, which is not really part of the control scheme (Pruszynski and Scott 2012; Scott 2004; Shadmehr and Krakauer 2008; Todorov 2004; Todorov and Jordan 2002). The loss function not only incorporates spatial constraints throughout the movement, but also many other factors like energy expenditure, movement time restrictions, reward contingencies, and so on (Diedrichsen 2007; Diedrichsen et al. 2009; Harris and Wolpert 1998; Izawa et al. 2008; Nagengast et al. 2010; Scott 2004; Serrancolí et al. 2017; Shadmehr and Krakauer 2008; Todorov and Jordan 2002).

While the loss function does not take center stage in most control schemes, here we present it as the interface between a non-motor and a motor stages. Specifically, we propose that the weight of each component of the loss function is determined by cognitive decision-making. Furthermore, we propose that loss function weights in the non-motor stage are adapted as a function of performance error, while the motor stage is directly influenced by the sensory prediction-error. In this framework, learning at the cognitive non-motor stage can alter behaviour without any downstream learning in the motor stage. The weights of the different elements of the loss function are thus specified as a function of cognitive action selection (decision-making) and can be updated on a trial-to-trial basis (Fig. 3) to take into account performance error (van Beers 2009, 2012) or a change in task requirement (Nashed et al. 2012; Orban de Xivry 2013; Orban de Xivry and Lefèvre 2016).

**Figure 3.**
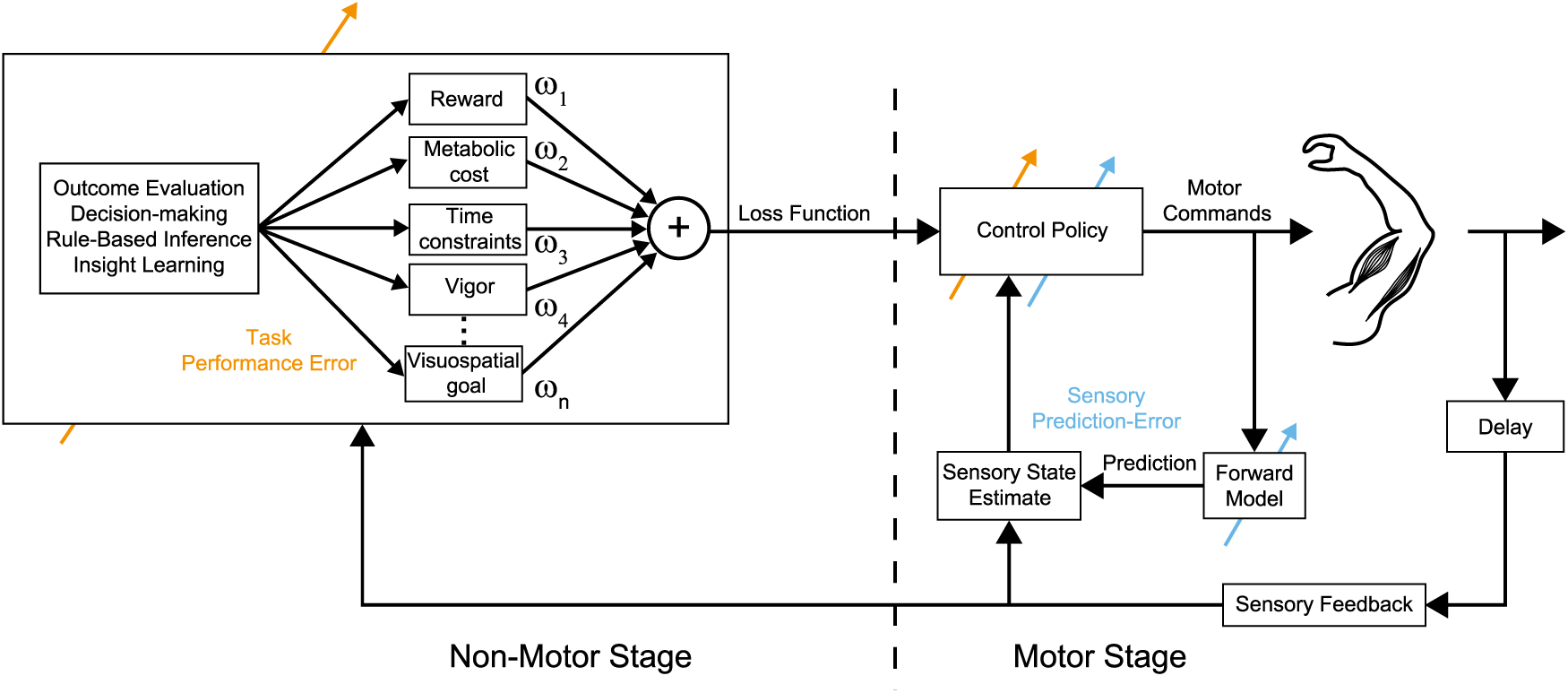
Schematic model for generating goal directed reaching movements and the locus of learning in visuomotor adaptation tasks. The control diagram is composed of non-motor and a motor stages. The motor stage contains all the elements of the conventional motor control scheme: it interacts with the environment via a body that is controlled by motor commands. Noisy sensory information about the state of the body is fed back to the motor system with a delay. This sensory information is integrated with the output of a forward model that uses an efference copy of the motor commands to predict the current and future state of the body. The motor commands are determined by the control policy (sometimes refer to as inverse model), which takes the loss function and the state estimate of the body as input and generates motor commands as an output. Optimal feedback control proposes that the motor commands result from the minimization of the loss function. In our schema, the loss function represents the interface between the motor and non-motor stages. The loss function is shaped by the non-motor part of the system in function of task requirements (instructed speed, accuracy demands, etc.) and internal factors such as motivation or vigor. These factors are not only sensitive to verbal instructions and apparent task conditions but also “high level” cognitive insight. Plasticity in this schematic model is represented by the colored arrows. The task performance error is computed at the level of the non-motor stage based on sensory feedback while the sensory prediction-error is computed in the motor stage based on the output of the forward model and the sensory feedback. We hypothesized that the sensory prediction-error could update both the forward model and control policy. Performance error primarily shapes the loss function but may also directly update the control policy.

We believe the operation of the non-motor stage requires further study in the context of adaptation tasks, but its general operation can be understood from research in other domains. Human decision making is chiefly concerned with explicit goals and rewards, here discussed as elements of a performance error feature vector. Human reasoning is the most flexible and insightful in the animal kingdom, sometimes resulting in abrupt “changes of mind” that emerge from re-evaluation of whether the current movement plan has the potential for success (Pacheco et al. 2020; Sharpe et al. 2019; Song and Nakayama 2009). For these changes of mind to occur, the predicted benefit of changing strategies must outweigh the costs in mental effort required to alter the established response contingencies (Kessler et al. 2009; Kiesel et al. 2010; McDougle and Taylor 2019; Monsell 2003; Orban de Xivry 2013; Orban de Xivry and Lefèvre 2016; Rogers and Monsell 1995). The idea that changes to the movement strategy rely on executive decision-making mechanisms is further bolstered by work that has consistently linking this learning to working memory capacity (Anguera et al. 2010, 2011; Christou et al. 2016; Vandevoorde and Orban de Xivry 2020a).

Once the loss function is specified at the non-motor stage, it is carried over to the motor stage where the control policy then generates a motor command (Fig. 3). As in other frameworks, this motor command is sent to the plant (spinal cord and body) to carry out the movement. A corollary discharge of this efferent motor command is relayed to a forward model (Bridgeman 1995; Sommer and Wurtz 2002, 2008), which predicts the future state of the body from the current state and motor command. This future state estimate is useful for many things, but in the context of learning, its sensory prediction is compared to the actual sensory feedback to calculate a sensory prediction-error (Jordan and Rumelhart 1992; Kawato 1999; Wolpert et al. 1995; Wolpert and Miall 1996). This sensory prediction-error influences behavior by adapting internal models (control policy and/or forward model). It is also possible that the performance error might affect the controller directly. We cannot be sure about the locus of implicit adaptation that is driven by target hitting error, given the available data. Although these learning is implicit, we believe it is possible for it to occur within the non-motor or motor stages. Due to a similar lack of data on temporally labile adaptation, we cannot be sure of its locus or even its driving error signal.

## The many errors and learning processes of visuomotor adaptation

Our description of the different errors and processes demands a new description of adaptation that is the sum of a many processes which differ with respect to their driving signal (performance error, sensory prediction-error and target hitting error) and with respect to how they are affected by the passage of time. We propose the existence of four distinct processes, whose contribution to total adaptation will depend on the experimental context.

First, the explicit process is driven by performance error (red curve in Fig. 4A). Over the course of adaptation, the contribution of this process increases rapidly, but later decreases until reaching a steady state. Notably, during the washout period (where the perturbation is removed), the explicit process has a negative contribution to the total adaptation. This is because the explicit process takes changes its aim to compensate for the aftereffects of the other processes so that it can maintain good task performance (Yin and Wei 2020). This process is only weakly influenced by the short passage of time—it is a declarative process which should have the same forgetting curve as other declarative memories (Ebbinghaus 1913; Squire 2004). The second process (blue curve on Fig. 4A) is temporally stable implicit adaptation driven by sensory prediction-errors. Third (cyan curve on Fig. 4A) is made up of adaptation that is driven by target hitting error that is stable with respect to the passage of time (Kim et al. 2019; Leow et al. 2018). The last process (green curve on Fig. 4A) is composed of implicit adaptation that decays over a timescale of seconds (Alhussein et al. 2019; Hosseini et al. 2017; Joiner et al. 2017; Joiner and Smith 2008; Sing et al. 2009; Vandevoorde and Orban de Xivry 2019; Zhou et al. 2017). There is not currently any published data bearing on which error drives temporally labile implicit adaptation.

**Figure 4.**
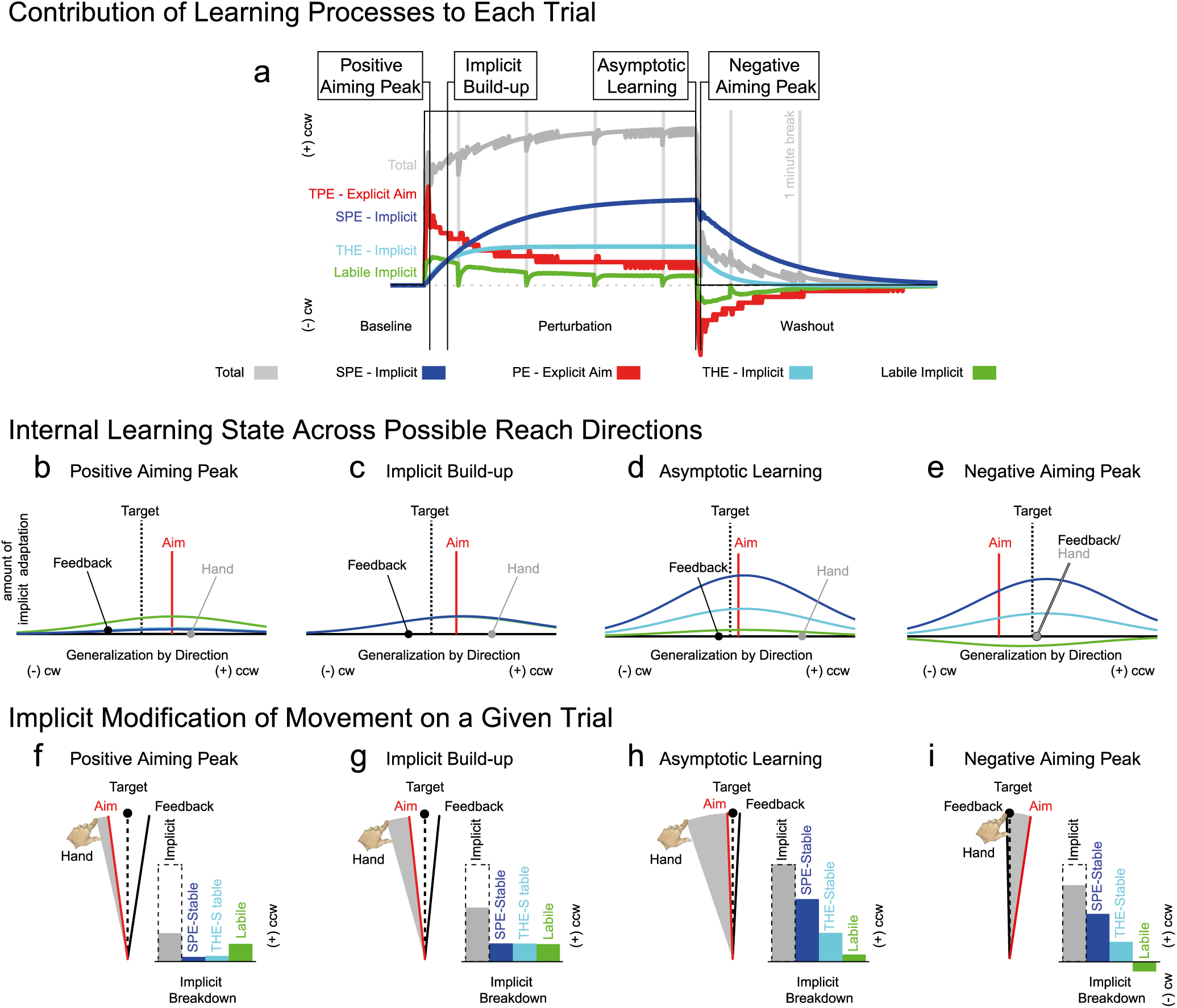
How the processes of visuomotor adaptation contribute to behavior during learning. A) Trial-by-trial evolution of the different processes during the adaptation period and washout in the presence of 1-minute breaks (grey vertical lines). Four important time points are highlighted and further described in panels B-I. The positive aiming peak corresponds to the trial where the aiming direction was furthest in counterclockwise direction. Implicit build-up corresponds to a time point early in perturbation block were implicit adaptation is small relative to the adjustment from explicit aim. Asymptotic learning corresponds to the end of the adaptation period where total adaptation is at its maximum. The negative aiming peak corresponds to the moment early in washout where the aiming direction is maximally clockwise. In this simulation, the participant aims at in the direction that will lead to best performance throughout the task. Note that the processes sum, although we have shown them each relative to zero. B-E) The internal generalization of implicit adaptation (SPE-implicit, THE-implicit and labile implicit) at the four time points highlighted above. The state of each process is represented across different possible movement directions (x-axis) relative to the visual target (vertical black dotted line). This shows the amount of implicit adaptation that is be predicted if the participant chose to aim for anywhere other than where they did. These curves show a peak of implicit generalization shifts with the point of explicit aim and that the implicit adaptation that is sampled on a given trial also depends on the explicit aim. Therefore, shifts in aim can cause a substantial change in the expressed amount implicit adaptation (change from D to E). F-I) Here, the effect of implicit adaptation (gray wedge) upon a given aimed reach is shown within the task workspace. The bar graphs show a breakdown of the total implicit adaptation (grey bars) into its constituent components (dark blue: SPE-implicit, cyan: THE-implicit, green: labile implicit). Notably, at the negative aiming peak, both the aim and labile learning are opposing the aftereffects caused by the temporally stable implicit adaptation.

These four learning processes we have described sum to affect behavior at any given moment. This causes indirect interactions between them, even though each process otherwise operates independently. For instance, learning from the SPE-driven stable implicit component of adaptation typically causes a decrease in performance error, even though this process is not driven by performance error. In some cases, the same learning process can actually increase performance error (Mazzoni and Krakauer 2006; Miyamoto et al. 2020; Taylor and Ivry 2011). Below, we describe how these processes evolve over the course of an adaptation experiment, and their hypothesized operation of each process in a variety of task contexts. It is our hope that this exercise provides an opportunity design future experiments that account for the operation of the processes we described above.

## Visuomotor Adaptation is the Sum of All Processes

In our framework, all the learning processes sum to influence the direction of a given reach. During baseline conditions, and at asymptotic learning, these processes reach a stable and predictable steady state, but their interaction can be complex and difficult to intuit at other points of the learning curve and especially near transitions in the experimental context such as the onset or offset of a perturbation (Fig. 4A). These interactions can be particularly counter-intuitive because implicit adaptation is generalized relative to the angle of aim (Fig 4B-E; Day et al. 2016; McDougle et al. 2017). This means that changes in aim during a visuomotor adaptation task necessarily shift the peak of implicit learning as the point of aim moves in the workspace (Fig. 4B,C). To point, if a participant aims far away from the target, the locus of implicit generalization will be far away from the target (Fig. 4B,E). This seemingly should have the biggest effect early in a rotation when aiming is greatest, but implicit adaptation has not yet had time to accumulate so it is of little consequence (Fig. 4B). As implicit adaptation grows, the difference between the peak of generalization at the aim point and implicit adaptation at the target will also increase (Fig. 4C). However, this implicit change in hand position will also cause explicit aiming to decrease so that good task performance is maintained (Taylor & Ivry, 2011; Taylor, Krakauer & Ivry, 2014; Miyamoto, Wang & Smith, 2020). This means that the peak of implicit generalization will be closest to the target when learning is asymptotic (Fig. 4D). Whether there will be much difference between the level of implicit learning at the target and at the point of aim depends upon the size of the perturbation, as larger rotations require aiming further from the target, and implicit adaptation is limited in the magnitude of its adjustment (Bond & Taylor, 2015; Morehead *et al*., 2017; Kim *et al*., 2017).

The effect of explicit aim on implicit learning levels is greatest when a perturbation is removed, or changes sign. In this case explicit aiming may change drastically (Fig. 4E), which will also cause the expressed level of implicit learning to change, and this may be a pronounced difference (Fig. 4I, negative aiming peak). This hopefully makes clear that aiming in visuomotor adaptation tasks is never “just aiming.” There is an underlying generalization of these processes, and they are related to the point of aim, as illustrated in Figure 4B-E. It is important to measure or control aiming throughout experiments because learning and expression of implicit motor memories are influenced by the explicit aim. The size and sign of these processes can vary depending on the conditions of the experiment, as shown by contrasting the simulated contribution of each process to behavior at specific trials in Figure 4F-I. Given the potential for many different combinations of these processes, it is important to understand how they operate and best to predict their operation with a model.

If aiming is not measured, or experimentally controlled, it can have important consequences for experiments. For instance, if two groups of participants showed similar learning during a perturbation, but aimed to different directions during washout, then the measured after-effects could be quite different (Fig. 5A-C). Whatever the reason for the difference in aiming during washout, a breakdown of these aftereffects into their constituent parts would reveal differences in both explicit and implicit learning processes. Without assessing aim, these aftereffects may be interpreted as implicit motor memories that whose spatial locus is the direction of the target. This is a mistake that has been made frequently with a prevalent experimental design that measures the decay of learning by removing visual feedback (Fig. 5A; Galea et al. 2011, 2015; van der Kooij et al. 2016; Shmuelof et al. 2012; Smeets et al. 2006). Many studies using this design did not disambiguate between implicit motor memories and explicit aiming strategies, complicating the interpretation of their data to the point that they need to be re-examined with these controls in place. To be specific, these studies have interpreted differences in aftereffects during no-feedback conditions as differences in implicit motor memory retention, when the differences are likely a function of aiming.

**Figure 5.**
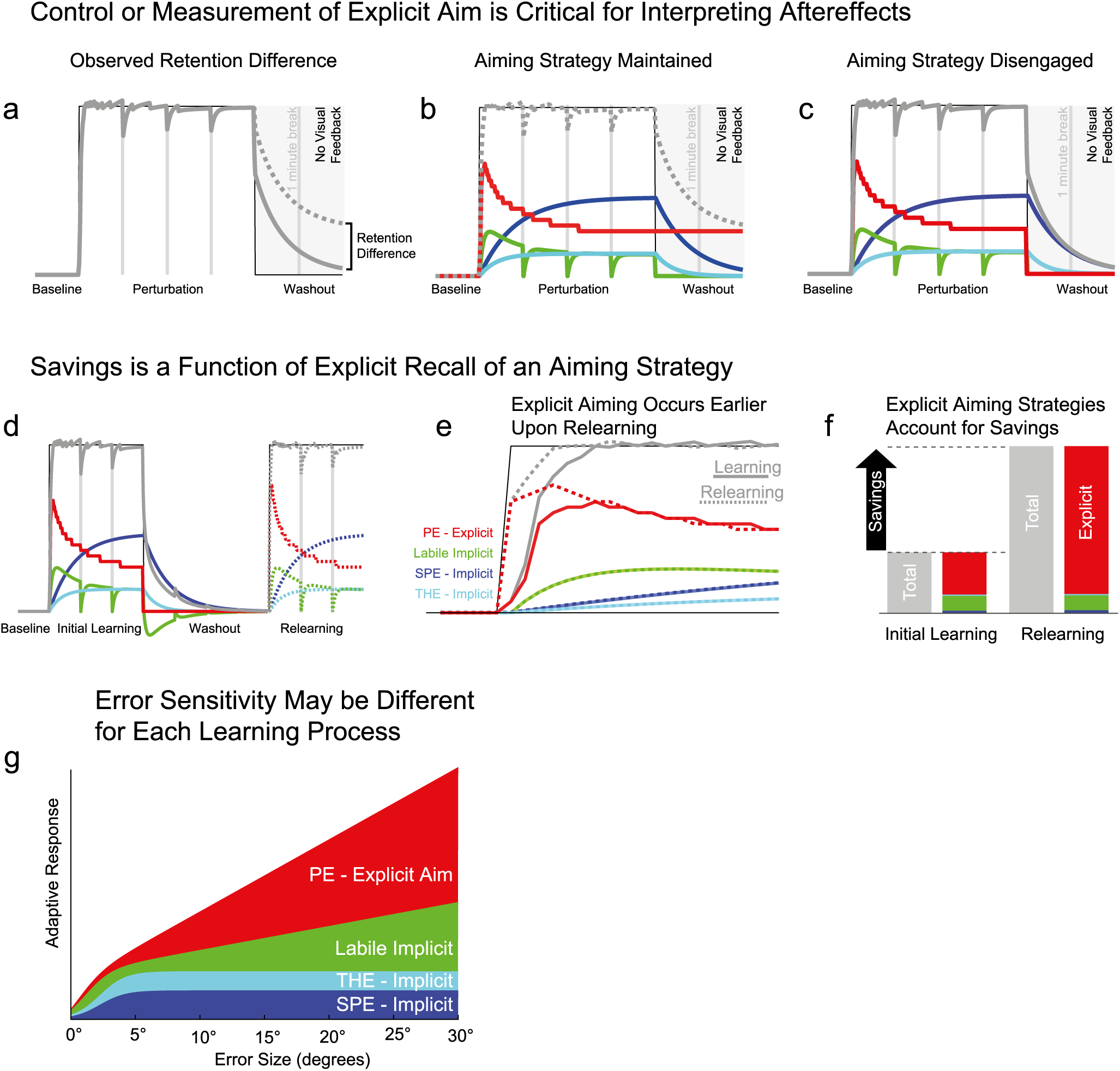
Specific cases where multiple learning processes can be misunderstood. First Row: control or measurement of explicit aim is critical for measuring an implicit aftereffect. A) Effect of instruction on total adaptation measured during a retention period (grey rectangle) in the absence of any visual feedback. The dotted curve illustrates differences in retention that have been observed in many experiments. B and C) illustrate how a simple change in aim can lead to the solid or dotted gray lines observed in A). Importantly, only aim has changed over these two simulations, but they can be misinterpreted as a difference in retention of an implicit motor memory if aim is not measured or controlled. Red, dark blue, cyan and green correspond, respectively, to the explicit, SPE-implicit, THE-implicit and labile implicit processes. SPE: sensory prediction-error; THE: target hitting error. Second Row: Savings upon relearning occurs because explicit aiming strategies are re-engaged during relearning. D) Evolution of the different processes underlying adaptation during learning and relearning. E) Same as D but learning and relearning are superimposed. Solid line: processes during learning; Dotted lines: processes during relearning. G) breakdown of total adaptation into the subprocesses for the first trials of the learning (left) and of the relearning (right) periods. Third Row: Hypothesized response for each component of motor adaptation at different error sizes. The x-axis provides the perturbation size while the y-axis reflects the amount of adaptation. The height of each layer represents the amount of adaptation from each of the processes. The sum of all the layers gives the total adaptation response to a specific error.

This complication for aftereffect assessment makes it necessary to use clear instructions for strategy use with these experimental designs. In experiments where the removal of the perturbation is cued and participants are instructed to give up any explicit strategies, behavior first shows an initial aim-related drop from instructions, and then subsequent changes in behavior directly correspond to the decay in implicit processes (Avraham et al. 2021; Codol et al. 2018; Hegele and Heuer 2010; Heuer and Hegele 2008; Holland et al. 2018; Taylor et al. 2014). Measurements of aftereffect magnitude and decay are best performed in the absence of feedback, as visual feedback could drive sensory prediction-errors, performance errors, or target hitting errors. In general, it is always a good idea to be cognizant of how errors will drive different learning processes.

A similar situation arises in savings designs, where the same perturbation is presented to the participants in two different experimental blocks separated by a period of rest or by a washout block. In this case, savings corresponds to a faster speed of relearning compared to the initial learning. Savings upon relearning has been repeatedly observed in visuomotor adaptation (Krakauer et al. 2005), but it is only recently that was this savings attributed to recall of an explicit aiming strategy (Haith et al. 2015; Morehead et al. 2015). In Figure 5E-F, we illustrate how savings arises solely from recall of an explicit strategy, with implicit mechanisms that learn at the same speed both initially and upon relearning. There remains some debate about whether implicit processes also show savings, but the lion’s share of the increase in relearning speed clearly arises from non-motor mechanisms and recent work suggests that implicit adaptation is actually attenuated upon relearning (Avraham et al. 2021; Huang et al. 2011; Leow et al. 2020; Orban de Xivry and Lefèvre 2015; Yin and Wei 2020).

So far, we have discussed block designs, where there is a strong incentive to change strategy when the same perturbation is used in a long set of trials (Morehead et al. 2015; Taylor et al. 2014; Wilterson and Taylor 2019). However, there is not necessarily utility in adjusting for an error when visuomotor perturbations are inconsistent or unpredictable (e.g. zero-mean random perturbation where perturbations of different amplitudes and direction are randomly interspersed among non-perturbed trials). The consistency of the perturbation across trials modulates the overall adaptive response (Fine and Thoroughman 2006, 2007; Gonzalez Castro et al. 2014; Herzfeld et al. 2014; Marko et al. 2012; Scheidt et al. 2001; Vandevoorde and Orban de Xivry 2019; Wei and Kording 2009). Decomposing this overall adaptive response into explicit and implicit mechanisms shows that the contribution of an explicit strategy increases with the predictability of the perturbation, while implicit adaptation is mostly invariant to the perturbation context (Avraham et al. 2019; Hutter and Taylor 2018).

We believe the operation of the motor and non-motor processes in visuomotor adaptation explain this difference in the response to errors when perturbations are consistent (on balance more likely for a given perturbation to occur) and inconsistent (or unpredictable) contexts. Explicit aiming strategies are chosen in the non-motor stage based on the expected utility of each movement strategy’s outcome (Skinner 1981; Trommershäuser et al. 2003, 2008). Therefore, when perturbations are consistent, the use of a strategy to counteract the perturbation will result in less performance error. However, if perturbations of different amplitude and direction are randomly interspersed, then aiming anywhere else will just as likely increase performance error as decrease it (Srimal et al. 2008). In this context it could take as little as a single trial for a participant to abandon an aiming strategy, as it is well established humans are remarkably sensitive the degradation in the contingency between the a particular movement strategy and its outcome (Fellows and Farah 2003; Shanks and Dickinson 1991).

This provides some clarity for the longstanding problem of sensitivity to error size in sensorimotor adaptation. Explicit strategies are selectively engaged if they are predicted to have a positive expected utility in terms of performance error (Fig. 5H). When employed, the contribution of the explicit component of adaptation to overall adaptation is roughly linearly proportional to the performance error size. Conversely, temporally stable implicit adaptation is marked by a linear response to error sizes below 5°, and a saturated response of the same size for larger errors (Avraham et al. 2019; Bond and Taylor 2015; Hutter and Taylor 2018; Kim et al. 2018). It is unclear whether temporally labile adaptation can be selectively disengaged in the manner of explicit strategies, but we have hypothesized that it is roughly linearly proportional to the composite error size. This error sensitivity function provides an outline of the trial-to-trial operation of the motor system in response to errors, when coupled with mechanism that will selectively engage explicit strategies depending on their expected utility.

Thinking about the motor system’s response to visuomotor perturbations as a composite of these processes also informs more complicated task designs, such as that employed by Mazzoni & Krakauer (2006). In this study, the researchers applied a visuomotor rotation of 45° but, after a few trials, informed their participants about the existence of the perturbation, its amplitude and its direction. In addition, a visual marker on the screen cued the aiming strategy that could be used to solve the visuomotor rotation. When instructed to do so, participants were able to bring their invisible hand towards the cued location in order to bring the cursor on the target in a single trial. While this experimental manipulation mostly negated the performance error, the sensory prediction-error was unchanged (the difference between the point of aim and the direction feedback was observed to move). This sensory prediction-error caused implicit adaptation in the direction opposite to the cursor rotation, which caused performance error to increase in opposition to the sensory prediction-error. This is a case where the interaction of explicit and implicit processes can be seen, so that explicit aim is adjusted in response to performance error even when strong instructions to aim at one specific point are provided (Taylor and Ivry 2011). The development of techniques to measure explicit aiming has helped clarify what occurs in this case, so that the overall behavior measured is best described as a dynamic tension of explicit and implicit processes that are driven by different error signals (Miyamoto et al. 2020; Taylor et al. 2014).

These examples in the context of aim-dependent generalization, after-effects, savings and error sensitivity demonstrate the importance of measuring the contributions of explicit and implicit processes or of controlling the errors that drive these processes in visuomotor adaptation experiments. It is important to anticipate the operation of these processes, and it ideally models should be used simulate how errors and learning will evolve during any given task.

## Looking beyond our current framework

We introduced a novel synthesis of the learning processes involved in visuomotor adaptation, and the signals that drive their operation. There are many implications of our framework, both proximally for sensorimotor adaptation experiments, and distally for general learning and skill acquisition. The fact that these processes exist and learn from different errors means that researchers must take care when designing experiments in humans. Moreover, this applies to animals where it is common to motivate participants with primary rewards during learning, and much more difficult to measure explicit awareness. Ideally, experiments would be designed so that the operation of each process and its driving signal is known. We believe this would greatly aid both the development of process-level models of sensorimotor adaptation and the understanding of its neurophysiological underpinnings. This approach, of breaking down the driving signals and constituent learning processes, can be generalized to other forms of learning such as sport or skill acquisition with brain-machine interfaces.

## Beyond visuomotor rotations

Our framework was developed from the results of visuomotor rotation experiments. However, we strongly believe that the errors and processes we outlined are operative in other forms of visuomotor adaptation. This includes gains applied to the radial component of a center-out reaches. In gain adaptation, the radial extent of the movement is perturbed by a given scale. For instance, making a 10cm point-to-point movement will be visually rendered as making a 12cm movement if the gain is set to 1.2. Humans adapt more quickly to gain adaptation than to visuomotor rotation and tend to generalize this learning globally to all reach directions, whereas learning from visuomotor rotations is generalized as mixture of local and global changes to other reach directions (Brayanov et al. 2012; Krakauer et al. 2000; Pine et al. 1996). Gain adaptation has not been explored in terms of explicit and implicit learning, or any of the other learning processes we have outlined. The learning differences between rotations and gains could come about may be because gain adaptation has a different balance of learning processes, (e.g. much more of an explicit component), or because the underlying mechanisms of implicit learning processes are fundamentally different (e.g. basal ganglia reward-dependent updating of global movement scaling versus sensory prediction-error updates of movement vectors). These possibilities for explicit mechanisms in the context of gain adaptation are so understudied that we cannot say almost nothing about them.

Similarly, we expect that our framework extends to prism adaptation, where the presence of multiple learning processes (Kornheiser 1976; Martin et al. 1996; Redding et al. 2005; Weiner et al. 1983; Welch 1969) and the importance of performance error and sensory prediction-error (Gaveau et al. 2014) have been acknowledged for a long time. Moreover, explicit aim has been measured throughout the course of prism adaptation (Leukel et al. 2015). We therefore expect that many of the processes we have outlined will play a role in prism adaptation, but it is also likely that there are additional processes that play a prominent role in prism adaptation. For instance, it is more common in prism adaptation for the subjective sense of straight ahead to be recalibrated, along with other senses like proprioception (Harris 1974; Redding et al. 2005). Our framework does not specifically account for recalibration of the senses, or high-level constructs like straight ahead because they are not observed in most contexts. Specifically, if the direction of movements is balanced as in a center-out reach task that span 360°, most of these effects will not occur. We believe it would be worth extending and modifying our framework to build a multiple process description of prism adaptation, although that is outside the scope of the current review.

Our framework is also relevant to another classic visuomotor perturbation, mirror-reflected visual feedback (Snoddy 1926; Starch 1910). It has been noted that the eventual learning from such feedback is more akin to *de novo* skill learning than adaptation (Hadjiosif et al. 2020; Lillicrap et al. 2013; Telgen et al. 2014). However, it appears that many, or all, of the processes we outlined in our schema are operative during this learning. Indeed, obligatory implicit adaptation that is driven by sensory prediction-errors may actually be maladaptive in such an environment, and such learning must be overcome with practice (Lillicrap et al. 2013; Wilterson and Taylor 2019). Learning at the level of explicit aiming can only partly explain this compensation, making it likely that the longer-term learning in mirror reversal tasks constitutes a skill learning process that is not outlined in our framework. How skill learning relates to our framework is pertinent question that we will address further below.

## Beyond perturbation of the visual feedback

The proposed schema was built upon results obtained in visuomotor rotation experiments, and it is easy to see how it generalizes to other forms of visuomotor learning. However, we believe that the errors and processes that we discuss and synthesize play a role in other forms of sensorimotor adaptation of reaching. These include adaptation observed in response to perturbations that affect intersegmental dynamics of the limb (Bresciani et al. 2002; Krakauer et al. 1999; Lackner and DiZio 1994; Sarlegna et al. 2010; Sarwary et al. 2013; Shadmehr and Mussa-Ivaldi 1994). There is good reason to think that neural locus of internal model adaptation for intersegmental dynamics is distinct from visuomotor adaptation, suggesting that there may be fundamental differences between adaptation of dynamics and visuomotor maps (Arce et al. 2010; Donchin et al. 2012; Perich et al. 2018; Rabe et al. 2009), but this does not necessarily mean that the types errors and learning processes that adapt movements are any different.

When adaptation of intersegmental dynamics is induced with force-fields, there is a clear contribution of both explicit and implicit learning processes (Hwang et al. 2006; McDougle et al. 2015; Schween et al. 2020). Explicit learning plays a smaller, but still substantial, role in the adaptation of dynamics compared to visuomotor adaptation. It is, however, difficult to compare the magnitude of these force field strategies with visuomotor contexts where the size of the perturbation determines the magnitude of strategy as it is both rare and complicated for force field experiments to compare adaptation to curl fields of different strength. Aside from the explicit/implicit divide, it has been shown that approximately ∼20% of the adaptation observed in single target force field studies is temporally labile (Sing et al. 2009). This limited information constitutes the extent of the available knowledge about the different processes and errors involved in adaptation of intersegmental dynamics, making it rather limited in comparison to visuomotor adaptation. We believe that it would be a great boon to understanding of motor learning if future dynamics adaptation tasks were designed to measure learning relative to performance errors, sensory prediction-errors, and the operation of the various implicit and explicit processes that we have outlined in our framework.

## Beyond perturbations of the upper limb

We expect that our framework should also generalize beyond the arm, to adaptation that occurs with any effector. For instance, in speech adaptation, participants speak words with specific phonemes and their auditory feedback is shifted in phoneme space, so that the auditory outcome is perceived as a different word (Houde and Jordan 1998). This distortion of the audiomotor mapping of speech is analogous to sensorimotor perturbations of the arm, and participants respond by changing their vocal output so that the auditory feedback matches that of their intended target word (Parrell et al. 2019). Performance error here is well-defined (difference between the target and produced sounds), and this performance error could engage explicit components of the speech production, and cognitive strategies similar to those seen in reaching, such as feedback corrections (Kim and Max 2021; Parrell et al. 2017). Similarly, speech adaptation is also sensitive to reward prediction-error (Parrell 2020). Speech adaptation is uniquely interesting because the movements involve control of the tongue, a which is a muscular hydrostat whose control involves no skeletal joint angles (Kier and Smith 1985).

In split-belt locomotor adaptation, people are asked to walk on a treadmill while the two belts of the treadmill move at a different speeds. Participants can compensate for this difference in speed by increasing the asymmetry in step frequency or length between the two legs (Dietz et al. 1994; Prokop et al. 1995; Reisman et al. 2005; Torres-Oviedo et al. 2011). Initially, it is common to see both compensatory mechanisms, but over training there tends to be a decrease in step length asymmetry using both spatial (change in the location of the foot landing) and temporal strategies (change in the timing of the foot landing) (Darmohray et al. 2019; Finley et al. 2015; Torres-Oviedo et al. 2011). A distinct, but similar, form of adaptation in step length asymmetry can be induced purely by perturbing visual feedback that corresponds to step length without changing belt speeds (Kim and Krebs 2012). Split-belt locomotor adaptation has been shown to have explicit components (Roemmich et al. 2016), and the gait adaptation induced by visual feedback alone can be attributed entirely explicit to an explicit strategy (French et al. 2018). This research suggests that a schema very similar ours should apply to locomotor adaptation. In the standard gait adaptation task where participants look straight ahead while trying to walk normally, it is difficult to define performance error as this is largely determined by the participant. For both the Roemmich and French studies, the addition of visual feedback for step length and the introduction of a goal state for this feedback helps with quantifying part of the performance error landscape, and it is possible to relate adaptive corrections to this quantity. It is important to note, however, that we should not assume that this artificial task performance feedback replaces more intrinsic elements of the performance error feature vector, such as a sense of stability, biomechanical or cognitive effort. Metabolic factors are likely given a much higher weighting in the loss function of locomotor adaptation, as the costs in walking are an order of magnitude greater than reaching (Finley et al. 2013; Huang et al. 2012; Sánchez et al. 2017). Given this reality, it has been suggested that energy minimization may be the primary driving factor in split-belt adaptation (Finley et al. 2013). Moreover, it is difficult to precisely characterize sensory prediction-errors, which may play a role in the implicit components of split-belt adaptation.

Eye movements can also adapt to external perturbations (Fukushima et al. 1996; Lisberger and Kahlon 1996; McLaughlin 1967; Paige and Sargent 1991). In saccade adaptation (Herman et al. 2013; Tian et al. 2009), the target that the participants directs his gaze to is displaced during the saccade so that vision of the target is suppressed. The target can be displaced along the primary axis of movement (gain up/down adaptation), or it perpendicular to the direction of movement (cross-axis adaptation). In line with our schema, saccade adaptation is well characterized by more than one learning process that adapt at different speeds (Chen-Harris et al. 2008; Ethier et al. 2008a, 2008b), and saccade adaptation can be driven (perhaps solely) by reward prediction-error as well (Madelain et al. 2011). Though there is a very salient performance error in this task, termed the retinal error, there is evidence that sensory prediction-error is one of the main contributors of saccade adaptation. Namely, if the eye lands between the expected landing position and the target, the sensory prediction-error points in the opposite direction than retinal error, the adaptation will follow the direction of the sensory prediction-error and not the performance error (Wong and Shelhamer 2010). Yet, it is clear that performance error does influence saccade adaptation. For instance, a clever manipulation that took advantage of the execution variability in saccades extinguished the target at the end of the saccade, but only for movements that were smaller than the median execution errors. This led to an increase in saccade amplitude, even though these movements should not have produced a sensory prediction-error (Collins and Wallman 2012). Thus far, studies have been limited to categorical eliminations of performance error or sensory prediction-errors. The parametric contribution of these errors to saccade adaptation have not yet been quantified.

Here we have discussed three types of sensorimotor adaptation that are commonly studied in humans, but there are many more. We believe that our general approach, characterizing errors and learning processes, can and should be extended to all these distinct forms of adaptation by experts in those subfields.

## Beyond motor adaptation

Our schema attempts to characterize the errors and learning processes that are involved in adaptation. An unresolved argument in the field of motor learning is whether sensorimotor adaptation is integral to, and representative of, motor skill learning, or whether it is a distinct form of learning that calibrates existing sensorimotor skills. Importantly, much of the theoretical basis of “error-based” learning and internal model theory was created to describe *de novo* learning of a multi-jointed robot arm, to only later be adapted to human motor learning (Atkeson 1989; Cheng and Sabes 2006; Jordan and Rumelhart 1992; Raibert 1978; Thoroughman and Shadmehr 2000). Models of sensorimotor adaptation are therefore models of motor skill learning at a fundamental level. Nonetheless, it has been argued that adaptation is a distinct learning process (Krakauer et al. 2019). We feel it is important to acknowledge this controversy, but wish to leave it aside, as we are of a mind that an application of our general aproach would greatly aid the study of skill acquisition throughout learning, whether or not it is one in the same as adaptation.

Skill learning has long been thought to involve more than one learning processes, and has long theorized that the total measured skill represents the summed output of subskills that are learned in parallel (Snoddy 1926; Starch 1910; Thurstone 1930). Moreover, the observation that skilled performance involves explicit and implicit mechanisms is nothing new (Adams 1987; Hendrickson and Schroeder 1941). Indeed, a primary characterization of skill learning is that performance transitions from an initial declarative process to a non-verbalizable, implicit state (Fitts and Posner 1967). Skill acquisition therefore has many parallels to our sensorimotor adaptation framework. What we believe is missing from this literature is a precise characterization of the signals, or errors, that drive this learning and, for any given skill, a description of the myriad processes that may be driven by these errors. As Thurstone (1930) described, each complex skill is likely composed of a set of subskills that learn at their own rates. To truly understand a complex skill, it is necessary to understand most of the component processes of the skill and what drives them, so that their evolution can be tracked and modelled over the course of learning.

## Beyond behavior

Many studies have attempted to elucidate the neural basis of sensorimotor adaptation. The methods employed are nearly as varied as the field of neuroscience itself, from the study of patients with neurological deficits to invasive electrodes implanted directly into the brain, non-invasive electrophysiology, transcranial magnetic and direct current stimulation, magnetencephalography, and magnetic resonance imaging of a functional, structural or spectroscopic nature. Unfortunately, all of the studies we could find were limited in light of our new framework because only few of them differentiated between the learning processes we outline (Jarosiewicz *et al*., 2008; Chase, Katz & Schwartz, 2012). Of primary importance, most studies did not measure or control the participants’ explicit intended movement. This means that the movement goal of the participant can vary substantially from one trial to the next without the experimenter’s knowledge. Instead of attributing the resulting changes in behavior to an explicit change in the intended movement, it is instead interpreted as arising from implicit learning or noise. This muddies the water tremendously, as implicit and explicit learning occurs in parallel and the brain areas that should be involved in these two types of processes should be quite different (McDougle et al. 2016; Taylor and Ivry 2012, 2014). This means that these studies will misclassify brain activity related to explicit strategy use. It is therefore no small wonder that brain areas commonly implicated in cognitive control and high-level decision making are often observed to be active during in sensorimotor adaptation (Anguera et al. 2010; Cassady et al. 2017; Ruitenberg et al. 2018; Seidler and Noll 2008; Shadmehr and Holcomb 1997).

Aside from the conflation of implicit and explicit processes, studies on the neural basis of sensorimotor adaptation face a substantial data analysis issue that we believe our framework can help resolve. Namely, the composite error, learning, and movement kinematics over the course of an adaptation experiment are all highly correlated with each other (Diedrichsen et al. 2005; Krakauer et al. 2004; Schlerf et al. 2012). We believe that the errors and processes we have outlined can be decoupled from one another, especially if model-based experiment designs are employed that maximize differences in the predicted operation of these processes. Some studies have started down this path with neurological patients, but many more studies using a variety of methods are needed (Butcher et al. 2017; Jahani et al. 2020; Morehead et al. 2017; Wong et al. 2019). As an example, post-movement event-related synchronization was first thought to be a function of (sensory) prediction-error (Tan et al. 2014) but later experiments with better control to differentiate between error types demonstrated that it was linked to performance error (experiment 2 in Torrecillos et al. 2015). We believe model-based designs that include errors and putative learning processes as regressors can help tremendously with these and similar issues.

## Extending behavioral methods beyond humans

As illustrated above, future research on the neural basis of sensorimotor adaptation would greatly benefit from a discrimination between the types of error and types of learning that we have outlined, both explicit and implicit. The methods that have been developed to parse these errors and processes were developed for use with humans, and many rely heavily on verbal instructions, precluding their direct use with animal models. This seems to present a substantial challenge, but we believe it is possible to translate many existing methods to animal models, such as non-human primates, monkeys, and rodents. Our suggestions here should apply to both experiments on visuomotor and force field adaptation.

First, it is important to note that explicit aiming is driven by performance error. Manipulations that affect the expected utility of a given movement strategy should therefore affect the use of explicit strategies. For animal experiments, the most promising method to manipulate performance error independently of sensory prediction-error may be that of Leow and colleagues (2018), where the target either jumps into or out of the path of the reach on every trial. This jumping target either always results in a miss or a hit, irrespective of the participant’s movement and sensory prediction-errors. This target jump method could be applied in the context of reaches or joystick tasks. In chimpanzees, a similar manipulation has been shown to modulate feedback corrections as a function of a visuomotor perturbation’s the task-relevance, although the focus of this study was not on sensorimotor learning (Kaneko and Tomonaga 2014). Similarly, the sensorimotor perturbation schedule itself can be manipulated, from fully inconsistent (zero-mean random) to fully consistent (blocked or alternating), where strategy use should scale from zero to full (Avraham et al. 2019; Hutter and Taylor 2018). Such task manipulations can be paired with delivery or withholding of rewards, or fully decoupled, but experimenters should carefully consider the performance error implications of direct rewards and punishments, especially on the strategic component of the task (Galea et al. 2015). Manipulation of the expected utility with the above methods can be combined with external sensory cues that are associated with various reward and/or perturbation conditions, as cue-action-outcome associations are easily acquired and are a reliable method for animals to induce a switch in behavioral strategies (Balleine and O’Doherty 2010; Sharpe et al. 2019; Skinner 1938). Conversely, external sensory cues have little or no capacity to modulate implicit sensory prediction-error driven learning over the short term (Cunningham and Welch 1994; Howard et al. 2013; Morehead et al. 2015; Vandevoorde and Orban de Xivry 2019, 2020a).

The preceding suggestions for animal experiments relate to the control of explicit strategies, but implicit adaptation can also be further broken down into subtypes. The measurement and characterization of temporally labile implicit adaptation is straightforward, as delays between trials will cause this learning to decay (Sing et al. 2009; Zhou et al. 2017). This learning can be measured precisely by the introduction of random delays of differing lengths, coupled with repeated retraining adaptation to asymptotic learning. Temporally stable learning can be dissociated into sensory prediction-error and intrinsic performance error adaptation by the target jump method of Leow *et al*. (2018). This method should induce greater implicit adaptation for enforced target miss over enforced target hit conditions. We believe that some clever experiments could result from the combination of these methods, or others, and we hope to see experimental designs employed in animal studies that discriminate between errors and learning processes that we have described for sensorimotor adaptation.

## Beyond traditional motor control

Brain-machine interfaces (BMI) are an alternative approach to study the neural control of movement. These techniques record neural activity directly and use it as an electronic control input (Fetz 1969; Vidal 1973) to interact directly with computers, robotic limbs or other devices (Carmena et al. 2003; Hochberg et al. 2006, 2012; Velliste et al. 2008). Here we focus on invasive BMI, where electrodes are implanted directly into the brain. This method is particularly elegant because the experimenter defines the relationship between neural activity and control signals, typically determining subsequent sensory feedback as well (Gilja et al. 2012). In a way, this is an advantage over traditional motor control, where the relationship between neural activity and movements is not fully understood. However, good control with BMI is not a given, as most systems require a substantial amount of training to achieve even mediocre performance (Farshchian et al. 2019; Ganguly and Carmena 2009; Orsborn et al. 2014; Pandarinath et al. 2017).

This extended training to enable BMI control means that it is fundamentally a sensorimotor learning problem. It also means that, like in other forms of skill learning, an approach that makes use of our schema could inform the theory behind its application. Performance errors in BMI control are very similar to visuomotor adaptation: this can be the difference between the intended and actual motion of the cursor, or it could be conveyed by points, sounds, or direct rewards and punishments, and so on. If our schema is anything to go by, performance errors may drive changes in the explicit aiming for BMI users. Interestingly, re-aiming has been a concern for some BMI researchers studying adaptation to decoder perturbations, and they have detailed how such a cognitive strategy would affect neural activity (Chase et al. 2012; Jarosiewicz et al. 2008; Zhou et al. 2019). Indeed, these non-human primate studies found clear evidence of re-aiming, which accounted for the majority of their learning in response to decoder perturbations. Human BMI studies also anecdotally report changes in movement strategy during learning: “The participant reported that… she was re-aiming to different directions to compensate for the applied perturbation…” (Sakellaridi et al. 2019). Importantly, although some BMI researchers take re-aiming and other cognitive strategies seriously, this is not the norm, and we believe that it should be.

Performance error drives explicit corrections, but BMI learning could be an ideal method to test whether it also drives implicit learning processes, as has generally been thought for skill learning. Indeed, it should be possible to contrast performance error with a putative sensory prediction-error in these tasks. It is as of yet unclear whether sensory predictions exist in BMI control, but their role in *de novo* learning has long been theorized, and these models should provide a rationale for detecting their presence or absence (Jordan and Rumelhart 1992). Temporally labile adaptation should also be fairly straightforward to test. Importantly, the advantage of invasive BMI control is that motor learning models could potentially predict the precise contribution of each neural unit to errors and model the role of each learning process in updating the activity of each neural unit.

## Beyond the current data: outlook

Our framework is built from visuomotor adaptation experiment results that have accumulated in recent years. While we believe this framework has explanatory power in many contexts, it also highlights a number of areas where the motor learning field is missing crucial information to fully understand motor adaptation. Some of these unknowns seem to perennially elude the field, while others are newfound questions that might be answered in a straightforward manner.

One particular issue relates to the specific types of internal models that are involved in sensorimotor adaptation. We have been careful to avoid attributing implicit learning in adaptation directly to a control policy (inverse model), or to a forward model. We believe that the currently available data cannot sufficiently rule out learning in either type of model, and it may be that adaptation arises from a modification of both types of model. We believe it is a real possibility that control policy and forward model modification are separate processes that learn at different rates, and potentially from different error signals. This is not altogether much different from the discussions around performance error and sensory prediction-error in Jordan & Rumelhart’s original 1992 paper about these error signals and learning processes. Similar to our uncertainty about the specific internal models driving implicit adaptation, we do not believe current data can specify which errors drive adaptation of inverse or forward models. This is mostly because the majority of past experiments have not differentiated between these error signals. These questions are of prime importance in moving the motor learning field forward.

Our framework incorporates several learning processes that we believe learn independently. Importantly, in asserting that these are independent, we mean that there are no documented interactions between these processes where the action of one process has unambiguously been shown to modulate that of another learning process. Rather, there is only evidence of indirect interactions, where one process changes factors like the locus of aim, or the amount of performance error, which in turn changes the inputs for a second learning process. In both cases, the actions of one process ultimately affects the behavior of another, but there is an important causal distinction between the two circumstances. We have detailed interactions of these processes in a limited number of contexts such as generalization, but there are many other ways that they could interact that depend on the task design.

It would be helpful to have a better understanding of the interactions between the adaptation processes we have outlined. For instance, it is clear that implicit learning, driven by a sensory prediction-error, can cause a performance error, which in turn drives explicit corrections (Mazzoni and Krakauer 2006; Miyamoto et al. 2020; Taylor and Ivry 2011). However, we know little from these studies about the role of target hitting error and temporally labile adaptation. We can model what these processes might do in the context of participants using a specifically provided aiming strategy, but it would be much better to have measurements of these processes in the midst of these complex tasks settings where implicit and explicit processes are in tension.

Similar to interactions between processes, it has been noted that the methods used to measure different adapted processes may lead to different results (Maresch et al. 2020b). We believe that more research in this vein is necessary, as subtle differences in experimental methods may also influence the processes we have outlined in different ways. For instance, the time required to give verbal instructions not to aim can cause the entire contribution of implicit temporally labile adaptation to decay. Although the intended experimental effect of these instructions is to eliminate the use of explicit strategies, its measured effect will also include the decay of implicit learning. Therefore, to avoid conflating these effects the verbal instruction group requires comparison to a control group that waits the same amount of time but receives no instructions to stop aiming.

Given the complexity of the interactions between learning processes, it is difficult to predict their behavior without a model. To point, we used models to make the learning schematics in this review and believe formal modelling work will be very helpful for designing experiments to measure these processes. It is, of course, crucial that this learning be modeled with the appropriate error signals, but it is equally important that these processes may learn from errors in different ways. Specifically, the internal learning mechanisms of explicit and implicit adaptation are likely quite different, but there may also be differences in how the implicit processes we outlined learn. Explicit learning should be broadly similar to other decision-making and reward learning models, using the performance error history to select from strategies based on their expected utility (Dayan and Daw 2008). Implicit learning is likely similar to state-space motor learning models (Cheng and Sabes 2006; Smith et al. 2006; Thoroughman and Shadmehr 2000), but with differences in error how errors map to corrections. For instance, the target hitting error from target hits and misses appears to be a binary signal (Kim et al. 2019) and sensitivity to sensory prediction-errors is non-monotonic (Kim et al. 2018).

In the past, modeling efforts for adaptation have primarily used the overall composite error as a unitary signal driving learning, even for models with more than one learning process (Cheng and Sabes 2006; Smith et al. 2006; Tanaka et al. 2012; Thoroughman and Shadmehr 2000). We believe this has obscured interesting questions of how performance errors, sensory prediction-errors, and intrinsic performance errors evolve over time. As discussed previously, a target hit may result in the elimination of both performance error and intrinsic performance error, but a putative sensory prediction-error remains, which will still drive adaptation (Mazzoni and Krakauer 2006; Shadmehr et al. 2010; Taylor and Ivry 2011). It is unclear how sensory prediction-errors evolve over the course of adaptation, despite the clear prediction from theory that it should be eliminated by forward model adaptation (Jordan and Rumelhart 1992). The paucity of experimental measures of sensory prediction-errors in sensorimotor adaptation is astounding, given its prominence in theoretical explanations of behavior. We believe studies focused on this and other errors could reveal a great deal about the operation of the motor system.

Model-based insights into the mechanisms of adaptation will not only be useful for process-level descriptions of this learning, but also for understanding its neurophysiological basis. Given the diversity of learning processes and errors, it is likely that sensorimotor adaptation recruits a wide number of brain regions. Although the cerebellum likely plays a prominent role in many facets of adaptation (both explicit and implicit) it is not the only brain area involved and is likely not the most important for some of these learning processes. Posterior Parietal, orbitofrontal, dorsolateral prefrontal and mediolateral prefrontal cortex may play a prominent role in explicit strategy use. These brain areas have been shown to be involved during sensorimotor adaptation (Diedrichsen et al. 2005; Shadmehr and Holcomb 1997; Taylor and Ivry 2014), but their specific attribution to explicit and implicit adaptation remains to be delineated. Although we believe both target hitting error-driven implicit adaptation and temporally labile adaptation are likely to be learning processes in the motor stage of our control diagram, we do not have any information to divine their specific neural locus. This leaves much room for exciting research on the neural basis of sensorimotor adaptation.

## Conclusion

We propose a wholistic framework for the understanding and interpretation of sensorimotor adaptation experiments. Our framework is primarily built from the last decade of observations made in the domain of visuomotor adaptation. This framework highlights the complexity of the myriad possible interactions between these processes during the course of typical and non-typical adaptation experiments. We view this complexity as an asset, rather than a liability. These errors and processes are distinct, and if care is taken, can be measured, modeled, and understood. The fact that these different errors and processes can be decorrelated presents a great opportunity to relate them to neural measurements. Visuomotor adaptation therefore features a tractable complexity, and there is still a great deal that we can come to understand about this form of learning. Importantly, as a field we are lucky that the cognitive strategies used in response to visuomotor rotations tend to be uniform and straightforward to measure. This simple fact opened the door to asking which errors drive which processes. By integrating these ideas with other observations, we have come to a new picture that explains several commonly observed effects in visuomotor adaptation experiments. It can and should be extended to explain even more.

Visuomotor adaptation can serve as a model for other forms of learning specifically because it has the right mix of simplicity and complexity. There are obvious parallels for other forms of sensorimotor adaptation, where the exact same learning processes may be operative, perhaps only differing in proportion. However, the motivating principles behind our framework can be applied more broadly to the acquisition of arbitrary skills from everyday life, such as darts, touch typing, billiards or tennis. By understanding the errors that serve as learning signals, and the processes that might learn from these errors, we may be able to move beyond simple learning curves to an understanding of the web of subskills contributing to a complex skill. We hope that we have succeeded not only in stimulating further research on visuomotor adaptation, but also on the broader application of the general approach we employed in developing our new framework for visuomotor adaptation.

## Acknowledgements

We were inspired and influenced by many great conversations with colleagues, especially including Jordan Taylor, Samuel McDougle, Richard Ivry and Maurice Smith. This work was funded by KU Leuven C1 grant: C14/17/115, and grant NS105839 from the National Institutes of Health.

